# Closed-loop real-virtual interactions validate 3D model of social coordination in fish

**DOI:** 10.64898/2025.12.07.692829

**Authors:** Ramón Escobedo, Justine Reynaud, Renaud Bastien, Stéphane Sanchez, Clément Sire, Guy Theraulaz

## Abstract

Collective motion in animal groups arises from social interactions rules, yet uncovering these rules requires quantitative models grounded in real behavior. We developed a fully three-dimensional, data-driven model of pairwise interactions in the schooling fish *Hemigrammus rhodostomus*, reconstructing attraction, alignment from experiments with two real fish swimming freely in a hemispherical bowl. Simulations of this model quantitatively reproduced empirical distributions of speed, distance, and orientation. We then embedded the model into a closed-loop virtual reality system, allowing a real fish to interact in real time with a virtual conspecific whose movements were governed by the same interaction rules. This biohybrid setup revealed that the model captures key social interactions underlying coordinated swimming. Our results establish a robust validation framework linking data, models, and behavior, paving the way for hybrid biological-digital collectives.

## 1 Introduction

Collective motion in animal groups is an archetype of distributed coordination. Fish schools, bird flocks, and insect swarms turn, accelerate, and reconfigure without central control, suggesting that local interaction rules can generate coherent global patterns.^1^ The central question is how to identify those interaction rules and test whether they are able to reproduce the observed coordination in realistic settings.^2^ Decades of work show that collective motion can emerge from simple alignment, attraction, and repulsion rules, modulated by noise and density. Models of self-propelled particles reproduce transitions between disorder and ordered motion and recover milling, swarming, and polarized states when parameters vary in plausible ranges.^1, 2^ At the same time, empirical studies emphasize biological specificity. Individuals differ in responsiveness, perception is anisotropic, and boundaries shape trajectories. These factors complicate the mapping from model rules to measured behavior.^3, 4^ Fish, in particular, offer a tractable system to bridge models and data. High-resolution tracking allows reconstruction of how burst-and-coast swimmers turn in response to neighbors and walls, revealing distinct alignment and attraction components that vary with distance, bearing, and relative heading. Previous work has shown that such behavioral analyses provide a solid foundation for constructing realistic models of schooling dynamics.^5, 6, 7^

Virtual reality closes this loop by embedding animals in responsive, programmable environments.^8, 9, 10^ Immersive systems for freely moving animals project visual stimuli that update in real time with the subject’s motion, enabling experiments with high ecological validity and precise control over sensory input.^11, 12, 13^ In fish, VR has been used to probe neural and behavioral mechanisms of social perception and prediction (Fig. 1A). Adult zebrafish in head-fixed VR exhibit robust, naturalistic behaviors, while neural imaging reveals circuits sensitive to prediction error when expected visual flow mismatches actual sensory input.^14, 15^ Recent work also reverse-engineers pursuit and schooling control laws using closed-loop VR, showing that simple proportional-derivative rules on egocentric distance can explain target tracking across contexts.^16^ VR studies further show that reciprocal interactions are necessary for natural temporal coordination, which emerges as an alternating rhythm of bursts that enhances spatial responsiveness.^17^

**Figure 1:**
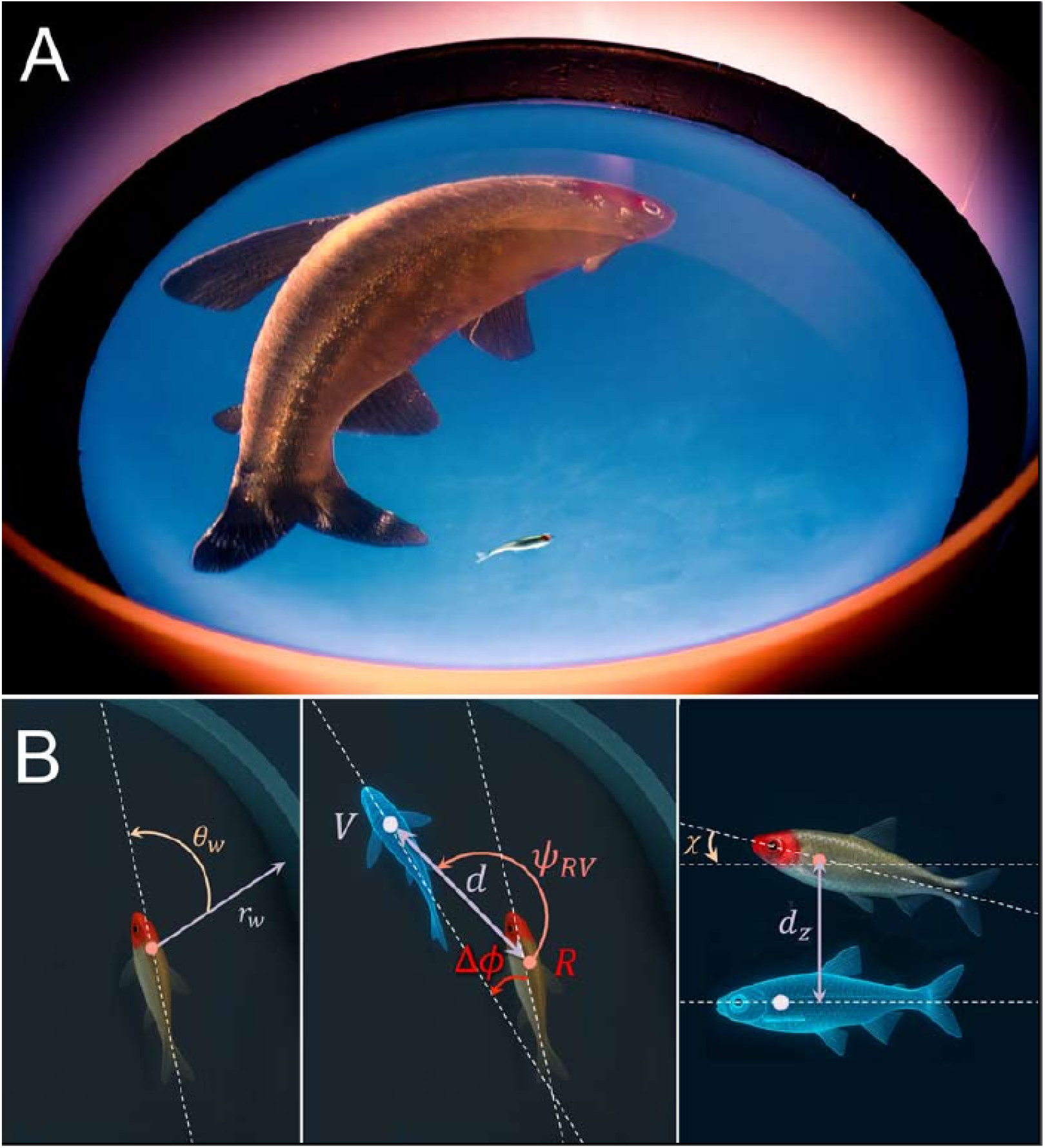
Closed-loop 3D virtual reality system and interaction geometry. (**A**) In the virtual reality setup, the 3D movements of a freely swimming fish are tracked in real time to feed a mathematical model of a virtual conspecific whose 3D anamorphic image is projected onto the inner surface of the hemispherical tank, allowing the model to continuously update the virtual fish’s position and orientation based on the real fish’s movements. (**B**) Variables used to describe the distance *r*_w_ and relative orientation of the real fish to the wall *θ*_w_ (left); the distance *d*, viewing angle *Ψ*_RV_ and relative orientation Δ*ϕ* of the real fish with respect to the virtual fish (middle), and the vertical angle of the real fish *χ* and the vertical distance between the real fish and the virtual fish *d*_*z*_ (left).

Despite this progress, a key gap remains. Most validations of collective-behavior models rely on statistical similarity between simulated and experimental distributions (see for instance^5, 6, 7^). Agreement in speed, spacing, or turning statistics is necessary, but not sufficient, to demonstrate that a model encodes the causal social interactions used by real animals. The stronger test is behavioral and consists in embedding a model-controlled agent into the interaction loop and asking whether a real conspecific responds as it would to a live partner. This test becomes more demanding in 3D, where vertical dynamics, wall effects, and the geometry of perception combine to shape motion. It also requires models that integrate individual self-propulsion, boundary interactions, and social responses into a single, data-driven framework.

Our study addresses this challenge in three steps, focusing on *Hemigrammus rhodostomus*, a small tropical freshwater species that exhibits remarkably cohesive schooling behavior and has become a model organism for studying the mechanisms underlying collective motion.^5, 18, 19, 6, 20, 21^ First, we quantify the interaction functions between two real fish swimming freely in a semispherical bowl. We reconstruct how horizontal acceleration decomposes into parallel and perpendicular components relative to heading, and how these components vary with velocity, neighbor distance, relative position, and heading difference, as well as with wall distance and incidence angle. We also characterize the vertical dynamics through vertical friction, preferred depth, and vertical attraction to a common depth. This reconstruction follows a data-driven approach that has proven effective in planar setups, extended here to full 3D kinematics.^22^ Second, we build a 3D self-propelled particle model that embeds those reconstructed functions, together with stochastic terms extracted from the data. We then test the model by comparing simulations of two virtual agents to experimental baselines of two real fish, assessing agreement across individual and collective observables. Third, we integrate the validated model into an immersive VR system.^23^ The anamorphic image of a digital twin of a fish is projected onto the inner surface of the bowl and controlled in real time by the model, using the tracked position and heading of the live fish as input (Fig. 1A). This hybrid setup lets us test whether the same interaction functions that reproduce real–real pairs in silico can also elicit natural coordination when one partner is virtual.

This pipeline enables three scientific questions. First, can interaction functions inferred from real–real pairs replicate individual and pair-level behavior in 3D simulations? Second, when one agent is virtual and controlled by the same model, does a real fish coordinate with it as with a real conspecific, and how do any deviations illuminate missing mechanisms such as actuation limits or latency? Third, how do the virtual partner’s speed and reciprocity modulate coordination strength, spacing, and alignment? Speed is a key control parameter for cohesion and polarization in fish groups, with faster kinematics often enhancing alignment and reducing reaction delays in both fish and birds.^24, 25, 26^ Reciprocity also matters: theory and experiments show that influence and leadership in moving groups depend on mutual responsiveness and timing.^27, 28^ We designed the closed-loop experiments to probe these mechanisms directly.

This approach complements neural and control-law studies in VR with zebrafish by focusing on full-body 3D kinematics in freely swimming social fish and by combining model reconstruction, forward simulation, and closed-loop behavioral validation in a single experimental framework.^14, 15, 16^ It also builds on the demonstration that visual cues alone can sustain coordinated motion in social fish and that immersive VR can evoke ecologically meaningful responses in freely moving species.^29, 30, 31, 32, 12, 33^ By placing a digital twin into a natural feedback loop, we move beyond validation at the level of static distributions and test directly whether the inferred social interactions produce the expected behavioral responses.

In sum, we address a core challenge in collective behavior and move from models that fit statistics to models that pass behavioral tests in closed loop. This hybrid strategy advances the validation of interaction rules in living systems and opens a path toward mixed collectives where biological and digital agents coordinate under shared rules.

## 2 Results

### 2.1 Quantitative analysis of pairwise swimming dynamics

Fig. 1B illustrates the main kinematic and geometric quantities used throughout this study to characterize and quantify fish motion, as well as the influence of social interactions on it. A detailed definition and mathematical description of these variables are provided in the Materials and Methods section.

Figure 2 shows the experimental distributions obtained from two real fish, together with their counterparts from numerical simulations of the model described below, for a set of key descriptors of individual and collective motion. The probability density functions of the full 3D speed v^3D^ and inter-individual 3D distance *d*^3D^ are nearly indistinguishable from those of their horizontal projections v and *d*, respectively (see SI Appendix, Fig. S1), indicating that motion remains mostly planar.

**Figure 2:**
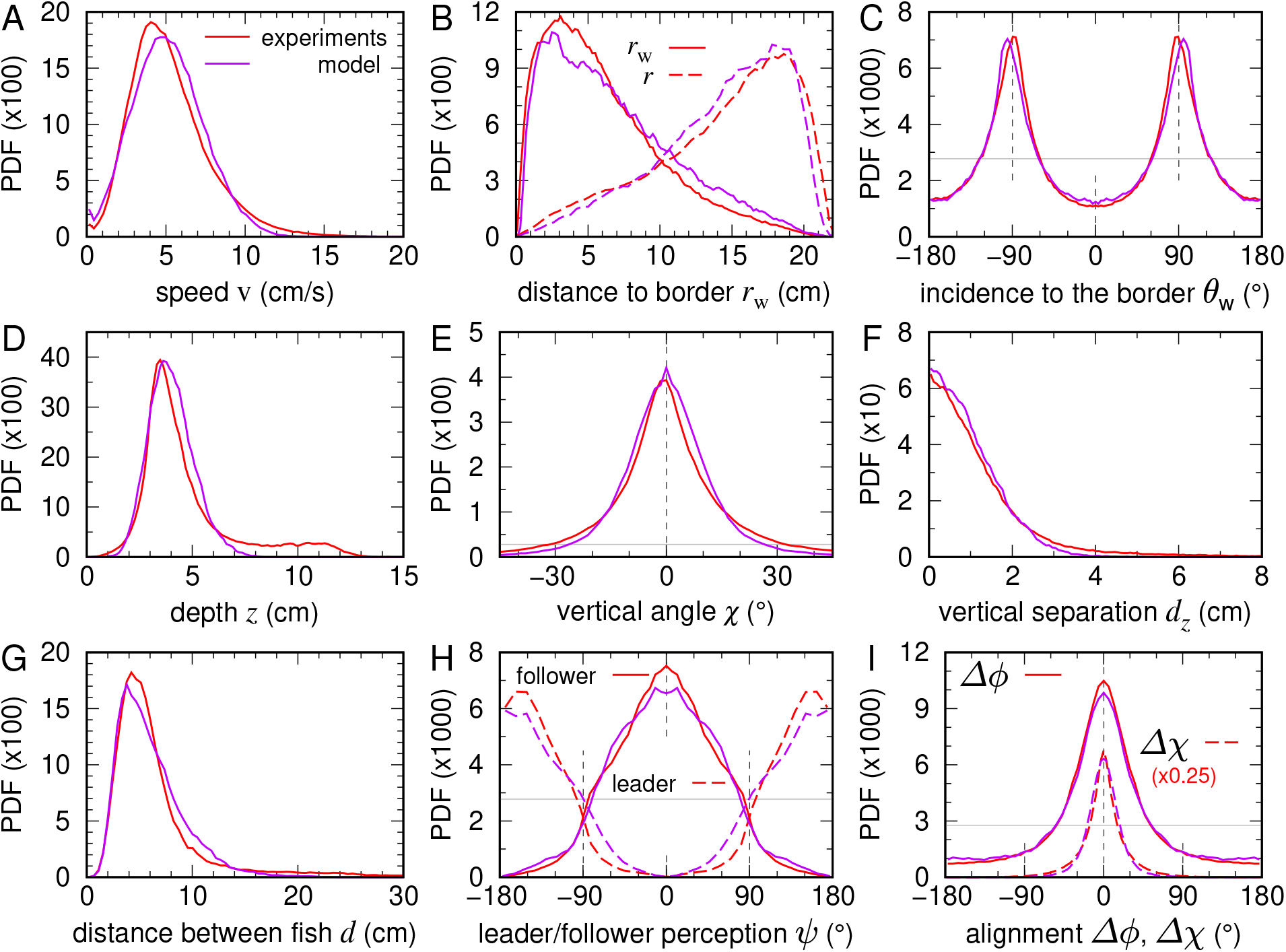
Quantification of free swimming behavior in a fish pair within the hemispherical bowl. Probability density functions (PDFs) of (**A**) swimming speed v, (**B**) distance to the bowl wall *r*_w_ (solid lines) and radial position *r* (dashed lines), (**C**) incidence angle relative to the wall *θ*_w_, (**D**) swimming depth *z*, (**E**) elevation angle *χ*, (**F**) vertical inter-individual distance *d*_*z*_, (**G**) horizontal inter-individual distance *d*, (**H**) viewing angle of the neighbor *Ψ* when acting as the geometrical follower (solid lines) or geometrical leader (dashed lines), and (**I**) heading misalignment in the horizontal plane Δ*ϕ* (solid lines) and vertical plane Δ*χ* (dashed lines). Red curves represent experimental data from two real fish, whereas purple curves correspond to numerical simulations of the model. Horizontal gray lines in (**C, E, H, I**) indicate the uniform angle distribution value, 1/360 (≈ 2.78 × 10^−3^).

Fish swam at a preferred speed of approximately v ≈ 5 cm/s (Fig. 2A), rarely exceeding 10 cm/s. Note that this speed range is markedly lower than observed in quasi-2D experiments in shallow water,^5^ where the mean speed was close to 10 cm/s, which might be ascribed to lower light levels in the VR experimental setup. The fish remained close to the wall of the bowl, with a mean wall distance *r*_w_ ≈ 3 cm (Fig. 2B), corresponding to a radial position *r* ≈ 18 cm from the central vertical axis. Their heading angle relative to the wall, *θ*_w_, was concentrated slightly below 90^°^ (Fig. 2C), indicating motion nearly tangential to the boundary. The fish also exhibited a preferred swimming depth around *z* ≈ 4 cm (Fig. 2D), with occasional excursions toward the bottom (uniform tail over *z* ∈ [7, 12] cm, with maxima up to 13 cm). Vertical motion was minimal overall: the elevation angle *χ* remained tightly centered around zero (Fig. 2E), confirming the strong preference for horizontal navigation.

At the collective level, individuals stayed in close proximity, with a typical horizontal separation *d* ≈ 5 cm (Fig. 2G) and a very small vertical separation *d*_*z*_ < 2 cm (Fig. 2F and SI Appendix, Fig. S1). As shown in Fig. 2H, a clear leader–follower configuration emerged: the follower typically viewed the leader directly ahead (solid curves peaked near zero), whereas the leader perceived the follower behind, with peaks near ± 160^°^ (dashed curves). Alignment was strong in both planes, as evidenced by the peaked distributions of heading differences Δ*ϕ* and Δ*χ* around zero (Fig. 2I), the latter being particularly narrow due to minimal vertical excursions.

Another notable behavior of the fish is their spontaneous tendency to perform U-turns while swimming side by side. One fish suddenly turns around to swim in the opposite direction of its companion, which in turn performs a U-turn to rejoin the first. SI Appendix, Fig. S4 shows the probability density function (PDF) and the survival curve of the time intervals separating two successive U-turns, displaying an exponential decay.

### 2.2 Data-driven 3D model of pairwise coordination in fish

Building on our empirical results and extending previous 2D modeling frameworks,^5, 6, 34, 7^ we develop a fully data-driven 3D dynamical model that accounts for the geometry of the hemispherical bowl and the social interactions underlying coordinated motion. Because of the limited precision of the three-dimensional tracking (especially along the vertical axis), segmentation of fish trajectories into discrete burst-and-coast phases was not feasible. As a consequence, we could not rely on the burst-and-coast–type description and model previously developed in two dimensions.^5^ It was therefore necessary to modify the model structure and adopt a continuous-time formulation of fish motion.

The instantaneous behavioral state of a fish is represented by the 9 variables v, v_*z*_, *r*_w_, *θ*_w_, *z, d, d*_*z*_, *Ψ*, Δ*ϕ*, respectively describing its horizontal and vertical speed, its distance and orientation with respect to the wall, its depth relative to the water surface, and its horizontal and vertical distance, viewing angle, and heading relative to another fish (see Fig. 1B). The temporal evolution of these variables combines biomechanical constraints, interactions with the environment, and social cues from the neighbor. The associated equations of motion are based on principles of self-organization, symmetry, and collision avoidance, and their functional forms are quantitatively inferred from the experimental data using a reconstruction procedure generalized from flat-arena studies to a curved 3D geometry.^5^ A detailed description of the model is provided in Materials and Methods.

We decompose fish motion into horizontal and vertical components:

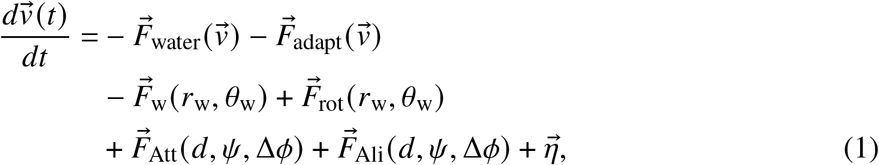

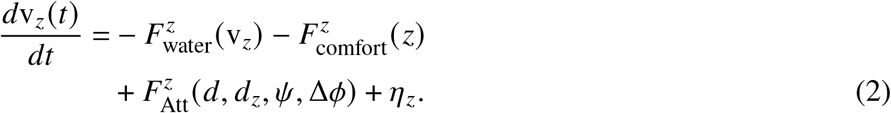

In the horizontal plane, the acceleration 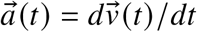 results from six components: **(i)** an effective friction emerging from hydrodynamic effects and the tendency of fish to swim at a preferred speed, 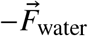, **(ii)** a term implementing the tendency of a fish to adapt its speed to that of its neighbor (or to the average speed 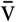 of its neighbors for more than one neighbor), of magnitude 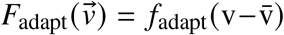, **(iii)** a radial repulsive interaction with the wall of the bowl, 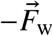, **(iv)** a wall-induced rotational force guiding motion tangentially, 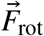, and **(v-vi)** the social attraction and alignment interactions with the neighbor, 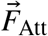 and 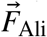 In the vertical direction, *a*_*z*_ (*t*) = *d*v_*z*_ (*t*)/*dt* arises from **(i)** a friction acting on the vertical speed 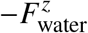, encapsulating the tendency of a fish to swim horizontally, **(ii)** a tendency to swim at a preferred comfort depth, 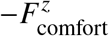, which also effectively results from the interaction of the fish with the wall of the bowl and the surface of the water, and **(iii)** a vertical attraction toward the neighbor’s depth, 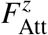. Additionally, a noise term is added to each component, 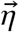 and 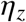 respectively, corresponding to the fish spontaneous decisions and random changes of heading.

A powerful assumption of our modeling process consists in considering that, once decomposed along the parallel and perpendicular components of the acceleration, the magnitudes of the forces are separable as products of single-variable functions, e.g., *F*_w_ (*r*_w_, *θ*_w_) = *f*_w_ (*r*_w_) *g*_w_ (*θ*_w_). Moreover, symmetry constraints apply to the angular functions to consistently reproduce the directional reactions of fish. For instance, the rotational force *F*_rot_ (*r*_w_, *θ*_w_) must be an odd function of *θ*_w_ at fixed *r*_w_ (hence, including a factor sin (*θ*_w_)) ensuring that a fish approaching the wall with *θ*_w_ > 0 (wall to its right) would turn in the opposite direction to that taken if it were approaching the wall with the opposite angle −*θ*_w_ < 0. This constraint preserves the right-left symmetry, which was shown to consistently hold for the considered fish species.^5^ Similarly, the alignment force *F*_Ali_ (*d, Ψ*, Δ*ϕ*) must include a factor sin (Δ*ϕ*) favoring a left turn when the neighbor is oriented counterclockwise with respect to the heading of the focal fish (Δ*ϕ* > 0) by increasing the perpendicular acceleration, while if the neighbor is oriented relatively clockwise (Δ*ϕ* < 0), the negative contribution induces a right turn.

Thus, with these separability and symmetry considerations, the magnitude of the forces induced by the wall is given by

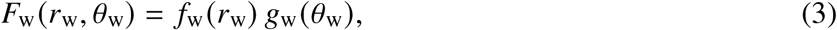

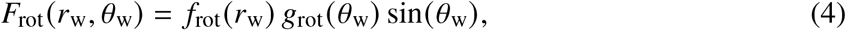

where *g*_w_ (*θ*_w_) and *g*_rot_ (*θ*_w_) are even functions of *θ*_w_. As for the social interactions, they can be written as

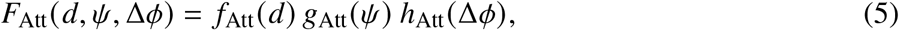

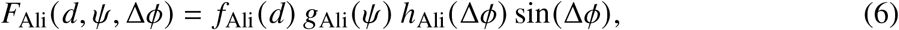

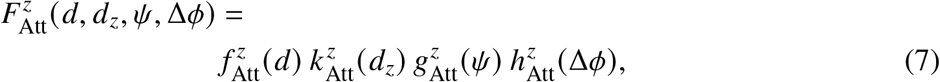

where all *g* and *h* functions of an angle are even. These interactions involve interaction functions depending on the distance between the fish and its neighbor or the wall, which are modulated by angular functions effectively describing the fish anisotropic perception of its environment.

The model also includes simple rejection rules that prevent the fish from crossing the physical wall of the bowl or the water surface. When a predicted step places the fish beyond these boundaries, the movement is rejected and a new fish orientation or vertical movement is recalculated (see Material and Methods for more details).

The resulting stochastic differential system is solved numerically using the Euler–Maruyama scheme with experimentally informed initial conditions.

### 2.3 Reconstruction of empirical interaction functions and validation of the 3D behavioral model

In this section, we present the interaction functions empirically reconstructed from the experimental trajectories and analyze their biological and dynamical roles in shaping coordinated motion. This method was originally introduced in^5^ and extended in^22^ and was extensively exploited to analyze the interactions arising from effectively 2D burst-and-coast trajectories, relevant to the case of fish swimming in shallow water. Here, we simply extend the method to 3D trajectories and to a continuous-time formulation of the equations of motion.

The general idea of the method consists in describing the different single-variable interaction functions introduced in the previous section by polynomial expansions, Fourier series, or tabulations. Ultimately, the parameters involved in this description are optimized to minimize the discrepancy between the observed acceleration of a real fish and the acceleration predicted by the model.

Some of the functions introduced above have a general shape that can be guessed on physical principles. For instance, the magnitudes (denoted with lowercase *f*) of the friction terms and the speed adaptation term are expected to resemble the following functional forms:

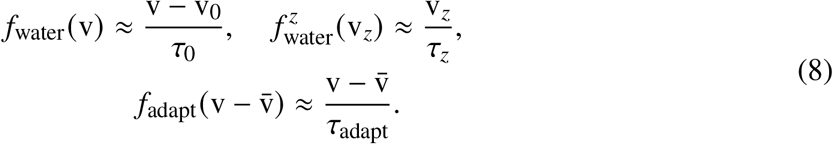

The friction *f*_water_(v) tends to maintain a speed v_0_ over a relaxation time *τ*_0_, while 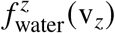 ensures that the mean vertical speed in a bounded bowl must necessarily be zero (i.e., that the fish is on average aligned with the horizontal plane). Finally, 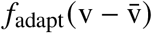 enforces the tendency of a fish to match its speed v to the speed 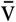 of its neighbor(s). In our reconstruction procedure, we are not imposing these simple but natural linear forms, and the functions are instead expanded as a finite order polynomial of their variable (in practice, we used third-order polynomials; see for instance, Eqs. (S1), (S2), and (S12) in Supplementary Materials). Similarly, some interaction functions are expected to vanish linearly for some value of their corresponding variable and will also be described by finite-order polynomials. This is the case for 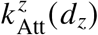 (expected to vanish at a vertical separation between fish *d*_*z*_ = 0) and 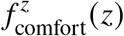 (expected to vanish at some preferred depth *z* = *z*_0_). Hence, in total, 5 functions are described as polynomials of their corresponding variable, whose coefficients are to be optimized by the reconstruction procedure.

Moreover, the model involves 8 angular functions (depending on the angles *θ*_w_, *Ψ*, or Δ*ϕ*) modulating the magnitude of the interaction of a fish with the wall and its attraction/alignment to its neighbor(s), and encoding the fish’s anisotropic perception of its environment. These angular functions are expressed as even Fourier series (preserving the left-right symmetry closely maintained by the considered fish^5^) of the form 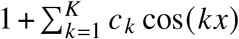, where the *c*_*k*_ ‘s are parameters to be optimized. We allowed up to *K* = 6 Fourier modes for each function, although for 2D trajectories, 2 or 3 modes were found to be sufficient to capture the different interaction functions.^5^

Finally, for the 5 interaction functions *f*_w_(*r*_w_), *f*_rot_(*r*_w_), *f*_Att_(*d*), *f*_Ali_(*d*), and 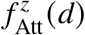, no general insight on their shape can be ascertained from physical/biological considerations. Hence, following,^5, 22^ we tabulate each of them over a finite grid (typically, 0.5 cm spacing for the grids for *r*_w_ and *d*) of their corresponding variable and consider the value of the function at each point of the grid as a parameter to be optimized. After optimization, the resulting functions are fitted by simple functional forms, which are ultimately implemented in the model. These fitting functions and the different polynomial and Fourier expansions introduced above are detailed in the Supplementary Materials.

The coefficients of the 5 polynomial expansions, those of the 8 Fourier series, and the parameters of the 5 tabulated functions are then optimized by minimizing the error between the experimentally measured accelerations and those predicted by the model. We use the total squared error Δ = Δ_∥_ + Δ_⊥_ + Δ_*z*_, where each term measures the squared difference between the experimental and model components of the acceleration along the fish heading, perpendicular to it, and along the vertical axis, *a*_∥_, *a*_⊥_, and *a*_*z*_, respectively (the detailed forms of *a*_∥_, *a*_⊥_, and *a*_*z*_ in the model are given in Materials and Methods). The matching error along the three directions are thus

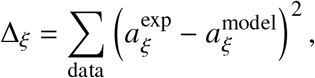

for *ξ* = ∥, ⊥, and *z*, and where each model acceleration is evaluated using the values of (v, v_*z*_, *r*_w_, *θ*_w_, *z, d, d*_*z*_, *Ψ*, Δ*ϕ*) observed in the corresponding experimental data.

Experimental accelerations are computed by finite differences after a slight smoothing of the experimental trajectories (see Materials and Methods). The model accelerations depend on the unknown functions introduced above. The coefficients describing their polynomial or Fourier expansion, or their tabulation parameters, are therefore the unknowns of the minimization problem. Because each force contributes linearly to the acceleration due to the separability assumption, Δ is quadratic with respect to the tabulated function values, thus allowing the efficient use of a gradient-descent optimization scheme to minimize the matching error Δ.

Figure 3 depicts the reconstructed functions (colored curves) superimposed on the PDF of their corresponding variable (gray areas). This highlights the regions of highest data density, where the reconstruction is most reliable and which primarily determine the overall shape of the functions.

**Figure 3:**
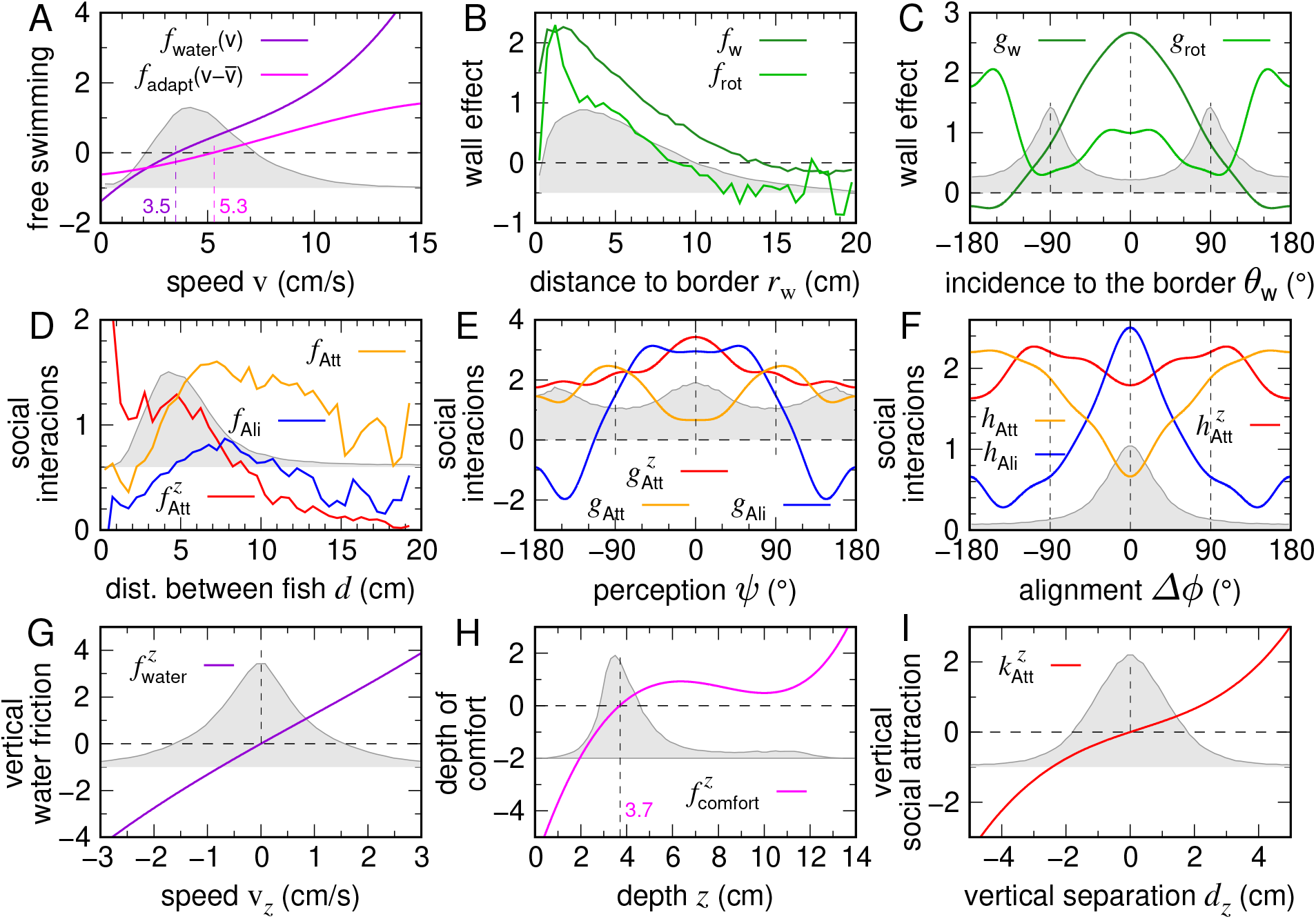
Experimentally reconstructed locomotion and interaction functions. Colored solid curves: reconstructed force components inferred from the experimental data. Gray areas: corresponding probability density functions (PDFs) of the associated state variables. (**A**) Self-propulsion terms: hydrodynamic friction and relaxation toward a preferred speed, *f*_water_(v) and tendency of a fish to adapt its speed to that of its neighbor, 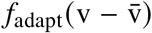 (arbitrarily centered at 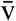 equal to the mean speed in this plot) as functions of the horizontal speed v. (**B**) Interaction with the wall: radial re pulsion *f*_w_ (*r*_w_) and wall-induced rotation *f*_rot_ (*r*_w_) as functions of the distance to the wall *r*_w_. (**C**) Angular modulation of wall effects through *g*_w_ (*θ*_w_) and *g*_rot_ (*θ*_w_) as functions of the incidence angle *θ*_w_. (**D**) Social interactions as functions of the horizontal distance between individuals *d*: horizontal attraction *f*_Att_(*d*), vertical attraction 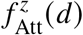, and horizontal alignment *f*_Ali_(*d*). (**E, F**) Angular modulation of the social interactions as functions of the viewing angle *Ψ* and heading difference Δ*ϕ*. (**G**) Vertical drag 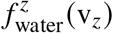 as a function of vertical speed v_*z*_. (**H**) Attraction toward a preferred depth *z*_0_ ≈ 3.7 cm, 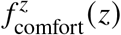, as a function of the depth *z*. (**I**) Vertical attraction between fish 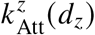 as a function of the vertical separation between fish *d*_*z*_. In these plots, the angular functions have been normalized such that 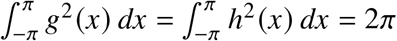.

The horizontal drag *f*_water_ (v) reduces speeds greater than v ≈ 3.5 cm/s, and increases speeds below it (Fig. 3A), while the speed adaptation function 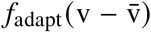 is essentially linear in the difference between the speed of a focal fish and that of its neighbor, as anticipated above (see Eq. (8)).

The wall-repulsion function *f*_w_ (*r*_w_) is strong when the fish is within 6 cm of the wall (i.e., about 2 body lengths, consistent with the results of^5^ for quasi-2D experimental setups) and essentially vanishes for *r*_w_ ≳ 13 cm (see Fig. 3B). The wall-induced rotational component *f*_rot_ (*r*_w_) has a similar spatial profile but a weaker magnitude. Both are modulated by even angular functions (see Fig. 3C) encapsulating the fish’s anisotropic perception of the wall, with the modulation of the repulsion to the wall *g*_w_ (*θ*_w_) peaking at *θ*_w_ = 0, when the fish is facing the wall, and being suppressed by a factor of 2 or more for |*θ*_w_| > 90^°^.

The horizontal attraction and alignment interactions *f*_Att_ (*d*) and *f*_Ali_ (*d*) initially grow with the distance between fish and then peak at *d* ≈ 2–3 body lengths (7–9 cm), before slowly decaying for *d* ≳ 12 cm, although the reliability of this long-range behavior is poor due to a lack of observed data, as fish are rarely found at such distances (Fig. 3D). The vertical attraction 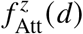 extends over a slightly narrower range.

Angular modulations reveal that social attraction is stronger when the neighbor is located laterally (maxima of *g*_Att_ near *Ψ* = ±90^°^, Fig. 3E), and more pronounced as the fish become more misaligned, with *h*_Att_ increasing as |Δ*ϕ*| goes from 0 to 180^°^ (Fig. 3F). Alignment is stronger when the neighbor is in front (|*Ψ*| < 60^°^, Fig. 3E) and becomes anti-alignment when the neighbor is behind (*g*_Ali_ < 0 when |*Ψ*| > 120^°^). Furthermore, alignment weakens as the fish are more misaligned (*h*_Ali_ decreases as |Δ*ϕ*| goes from 0 to 180^°^; see Fig. 3F).

As anticipated above (see Eq. (8)), the vertical friction 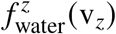 is a nearly linear odd function that suppresses vertical excursions (see Fig. 3G). The effective force 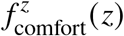 driving the fish toward a preferred depth *z*_0_ ≈ 3.7 cm (see Fig. 3H) exhibits a stronger/steeper repulsive interaction with the surface of the water (*z* = 0) than with the bottom of the bowl. For the interaction between fish along the vertical direction, the observed modulation by *Ψ* and Δ*ϕ* is weak, indicating that vertical attraction depends primarily on the depth difference (the weakly nonlinear and almost odd 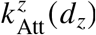 in Fig. 3I) and on the distance between fish 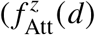 in Fig. 3D, which is strongly suppressed for *d* > 10 cm).

Figure 2 shows that the agreement between experimental data and simulations of the model integrating the empirically reconstructed interaction functions is excellent. This agreement holds for both horizontal kinematic variables, such as speed, relative position, and heading, and vertical components, including swimming depth *z* and vertical orientation *χ*. Moreover, the model also accurately reproduces the time intervals separating two consecutive U-turns (SI Appendix, Fig. S2).

### 2.4 Closed-loop experiments with a virtual conspecific controlled by the model

Having established the natural coordination patterns of two real fish and derived a data-driven model that accurately reproduces these dynamics, we next examined how a real fish behaves when it interacts with a virtual conspecific whose motion is controlled in real time by the model.

We tested three closed-loop conditions, each differing in the behavioral parameters assigned to the virtual fish:

- **Condition C1**: a perfect clone, reproducing the behavior of a real fish in a pair, swimming with a mean speed v ≈ 5 cm/s;
- **Condition C2**: a faster clone moving at twice the speed used in condition C1 (v ≈ 10 cm/s);
- **Condition C3**: a faster clone identical to the one used in condition C2 but which is independent of the real fish, creating a one-way interaction where only the real fish responds.

These closed-loop assays allow us to quantify how a real fish adjusts its behavior depending on the speed and responsiveness of the virtual partner. By comparing the resulting probability distributions of key behavioral measurements with those obtained for two real fish (Fig. 4), we can evaluate both the realism of the model and the extent to which coordination mechanisms remain engaged when interaction becomes kinematically challenging or asymmetric. Remarkably, despite substantial speed differences introduced in C2 and C3, and the absence of reciprocal interaction in C3, the real fish still exhibits similar social responses (i.e., similar pairwise separation and alignment) in response to the virtual conspecific.

**Figure 4:**
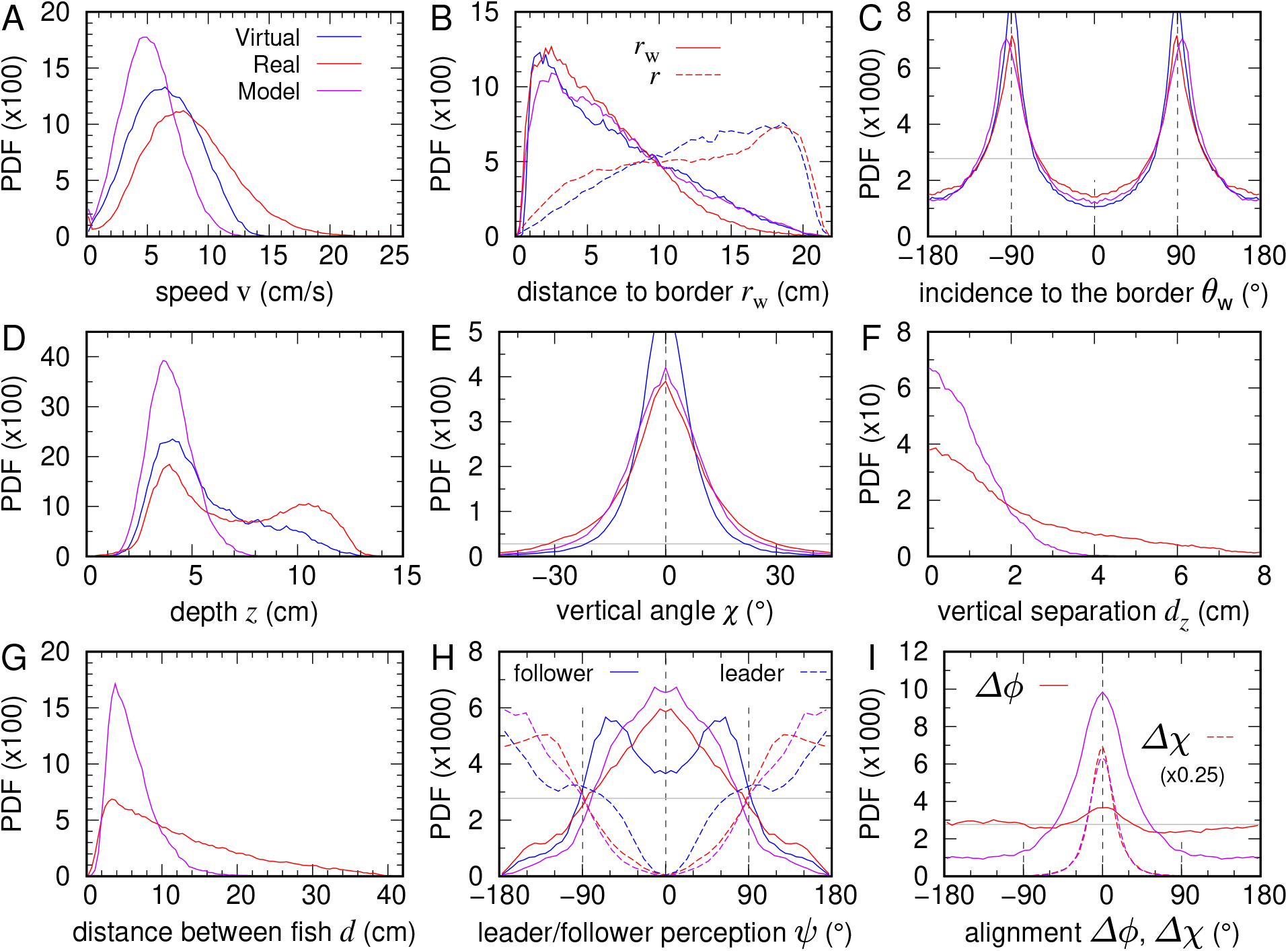
Model validation through real–virtual fish interactions. Probability density functions (PDFs) of behavioral variables measured for the real fish interacting with a virtual conspecific controlled in closed loop by the model (red), for the virtual fish (blue), and for numerical simulations involving two real fish (violet). (**A**) Swimming speed v. (**B**) Distance to the wall *r*_w_. (**C**) Incidence angle relative to the wall *θ*_w_. (**D**) Swimming depth *z*. (**E**) Elevation angle *χ*. (**F**) Vertical inter-individual distance *d*_*z*_. (**G**) Horizontal inter-individual distance *d*. (**H**) Viewing angle *Ψ* for geometrical leader (dashed lines) and follower (solid lines). (**I**) Heading alignment between the real and virtual fish in the horizontal plane Δ*ϕ* (solid lines) and in the vertical direction Δ*χ* (dashed lines).

#### 2.4.1 Behavioral responses of a real fish interacting with a realistic virtual partner

We refer to the condition where both fish are real as the 2R (two real) baseline. Figure 4 reports the same observables presented in Fig. 2 for condition 2R, but now corresponding to condition C1, where the real fish interacts with a model-controlled virtual clone.

The probability density functions for the real fish (red), the virtual fish (blue), and simulations of two real fish (violet) allow us to directly compare (i) the behavior of a real fish when paired with a real versus a virtual partner and (ii) how the model-defined behavior changes when the real fish deviates from its natural pairwise dynamics.

Overall, the qualitative features of bowl swimming remain present: real and virtual fish stay near the wall (Fig. 4B), swim nearly tangentially to it (Fig. 4C), maintain a preferred depth (Fig. 4D), and exhibit small vertical angles (Fig. 4E). Coordination is still visible, as shown in SI Appendix, Movie S1 and SI Appendix, Fig. S2.

However, substantial quantitative differences emerge. Most strikingly, the real fish swims markedly faster when interacting with the virtual clone than with a real conspecific: the PDF of its speed peaks near v ≈ 8 cm/s in C1, reaching up to 20 cm/s, whereas in 2R the speed rarely exceeds 10 cm/s (Fig. 4A). In fact, the swimming speed of the real fish now aligns with the characteristic speeds documented in quasi-2D experiments.^5^ As noted earlier, the projection of the virtual fish may partially restore lighting conditions comparable to those of the 2D setup, which could explain the observed agreement in swimming speeds. The virtual fish also swims faster than implemented in the model (peak near v ≈ 6.5 cm/s instead of 5 cm/s), thanks to the speed adaptation term (see Eq. (8) and the discussion below it). However, as it is constrained by the model’s speed parameters, it cannot accelerate sufficiently to keep up with the real fish (see SI Appendix, Movie S1). This speed mismatch produces an increase in the typical distance between fish (Fig. 4G) and prevents the virtual fish from positioning itself appropriately to maintain alignment (Fig. 4I).

A second major difference is a marked increase in bottom exploration (Fig. 4D). While the real fish in 2R mostly remains near its preferred depth *z* ≈ 4 cm, the real fish in C1 frequently visits the lower region of the bowl, producing a secondary PDF peak around *z* ≈ 10.5 cm. The virtual clone exhibits the same trend, though less strongly.

These changes weaken spatial cohesion: the PDF of the horizontal distance between fish *d* broadens substantially, with *d* > 20 cm observed frequently (Fig. 4G). Alignment is also drastically reduced: the distribution of Δ*ϕ* is almost flat (Fig. 4I), indicating minimal heading correlation.

The PDF of the angle of perception shows that fish still display the classic spatial configurations of geometric leader and follower. As a follower, the real fish predominantly perceives the virtual one ahead (|*Ψ*_follower_| < 90^°^; red solid line, Fig. 4H), consistent with 2R behavior (purple line), and as leader, it perceives its neighbor behind (|*Ψ*_leader_| > 90^°^; red dashed line, Fig. 4H). However, the virtual fish tends to perceive the real one laterally (|*Ψ*| ≈ 60^°^; blue solid line), revealing a breakdown in reciprocal leadership. As geometrical leaders, both fish maintain *Ψ* distributions similar to real-pair interactions. The virtual fish’s behavior differs subtly from that of the real one. As leader, its behavior remains similar to that observed in interactions between real fish (blue dashed line, Fig. 4H). However, as a follower, it clearly keeps the real fish in front but at one side, as shown by the two peaks of the PDF of *Ψ*_follower_ at ±60^°^ (blue solid line, Fig. 4H), thus breaking the reciprocity typically observed in pairwise interactions.

To investigate the origin of these deviations (faster motion, deeper swimming, larger separations, and reduced alignment) we quantified turning behavior and acceleration components.

The real fish exhibited much stronger turning dynamics when interacting with the virtual clone. SI Appendix Fig. S3A shows the PDF of the signed angular speed *dϕ*^+^/ *dt*, defined as the instantaneous heading change signed with the angle of incidence to the wall *θ*_w_, so that left and right turns can be accumulated. The PDF peaks at *dϕ*^+^ / *dt* ≈ 47.5^°^ s^−1^ in C1, compared with only ≈ 21^°^ s^−1^ in 2R, and shows a greatly increased probability of extreme turns (high-value tail for *dϕ*^+^ / *dt* > 100^°^ /s in purple line). We also note that the mean time separating a U-turn performed by one fish and that of its companion is higher in C1, 6.6 ± 12.7 s, than in the 2R condition, 3.2 ± 5.1 s (SI Appendix, Fig. S5). Since the time intervals between U-turns executed by either the real or the virtual fish’s partner are similar, this suggests that the virtual fish’s inability to keep up with the real fish stems from its limited acceleration capability.

Kinematic forces are also amplified. The absolute mean parallel acceleration nearly doubles (⟨|*a*_∥_ |⟩ ≈ 5.2 cm s^−2^ in C1 vs. 2.5 cm s^−2^ in 2R), and the perpendicular acceleration nearly triples (⟨|*a*_⊥_|⟩ ≈ 9.9 cm s^−2^ in C1 vs. 3.5 cm s^−2^; SI Appendix Fig. S3B,C).

Together, these observations reveal that although the model-controlled virtual fish can elicit coordinated interactions, the lack of full reciprocity and the inability of the virtual fish to match the real fish’s acceleration capabilities lead to a fundamental disruption of pair cohesion and alignment.

#### 2.4.2 Behavioral response of a real fish to a faster virtual conspecific, with or without reciprocal interactions

In Condition C1, the real fish swam considerably faster than when paired with a real conspecific in the VR setup. This observation motivated us to examine how a real fish responds when the virtual partner swims at that same higher speed, which coincides with the typical real fish speed observed in quasi-2D setups.^5^ We refer to this configuration as Condition C2. Furthermore, to test the role of reciprocal interaction, we implemented a third configuration (Condition C3) in which the virtual fish behaves as an isolated agent and does not respond to the real fish. Any coordination in C3 thus arises solely from the real fish adapting to the virtual one.

For each condition, we plotted the PDFs of the relevant behavioral variables: Condition C2 shown in the upper rows of Fig. 5, Fig. 6, and SI Appendix Fig. S6, and Condition C3 in the bottom rows. We also include numerical simulations in which both agents follow the behavior of the virtual fish. In C3, these simulations represent a “null model” without reciprocal interactions. The parameter values of the model used in these different conditions are provided in SI Appendix, Table S1.

**Figure 5:**
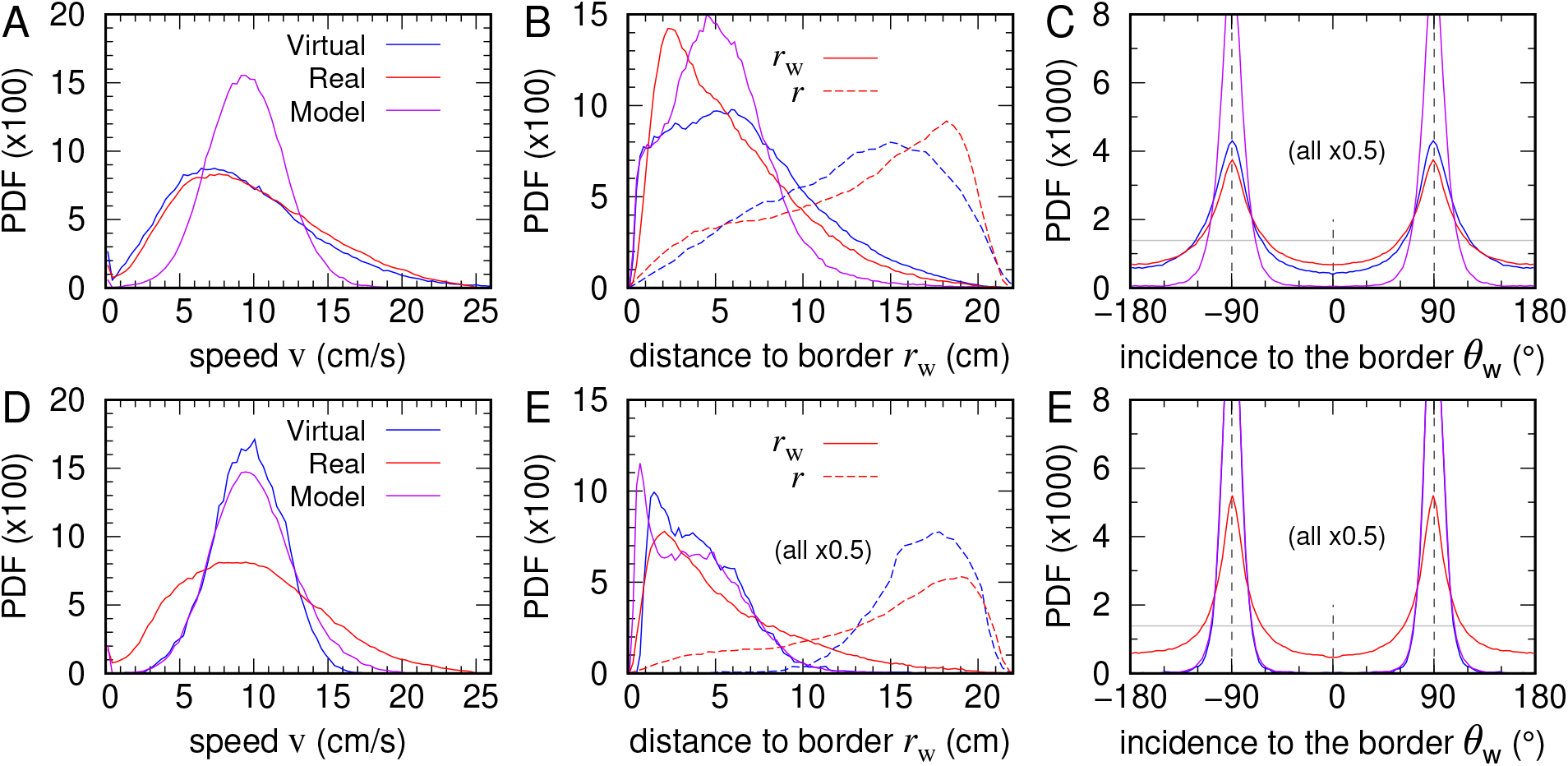
Individual behavioral responses to a faster virtual conspecific with and without social interactions. Probability density functions (PDFs) of real fish (red), the virtual fish (blue), and simulations of the 2R-based model (violet) at a higher mean speed v ≈ 10 cm/s. (**A–C**) Condition C2: with social interactions enabled (bidirectional coordination). (**D–F**) Condition C3: without social interactions (unilateral interaction). (**A**,**D**) Swimming speed v. (**B**,**E**) Distance to the wall *r*_w_. (**C**,**F**) Incidence angle relative to the wall *θ*_w_.

**Figure 6:**
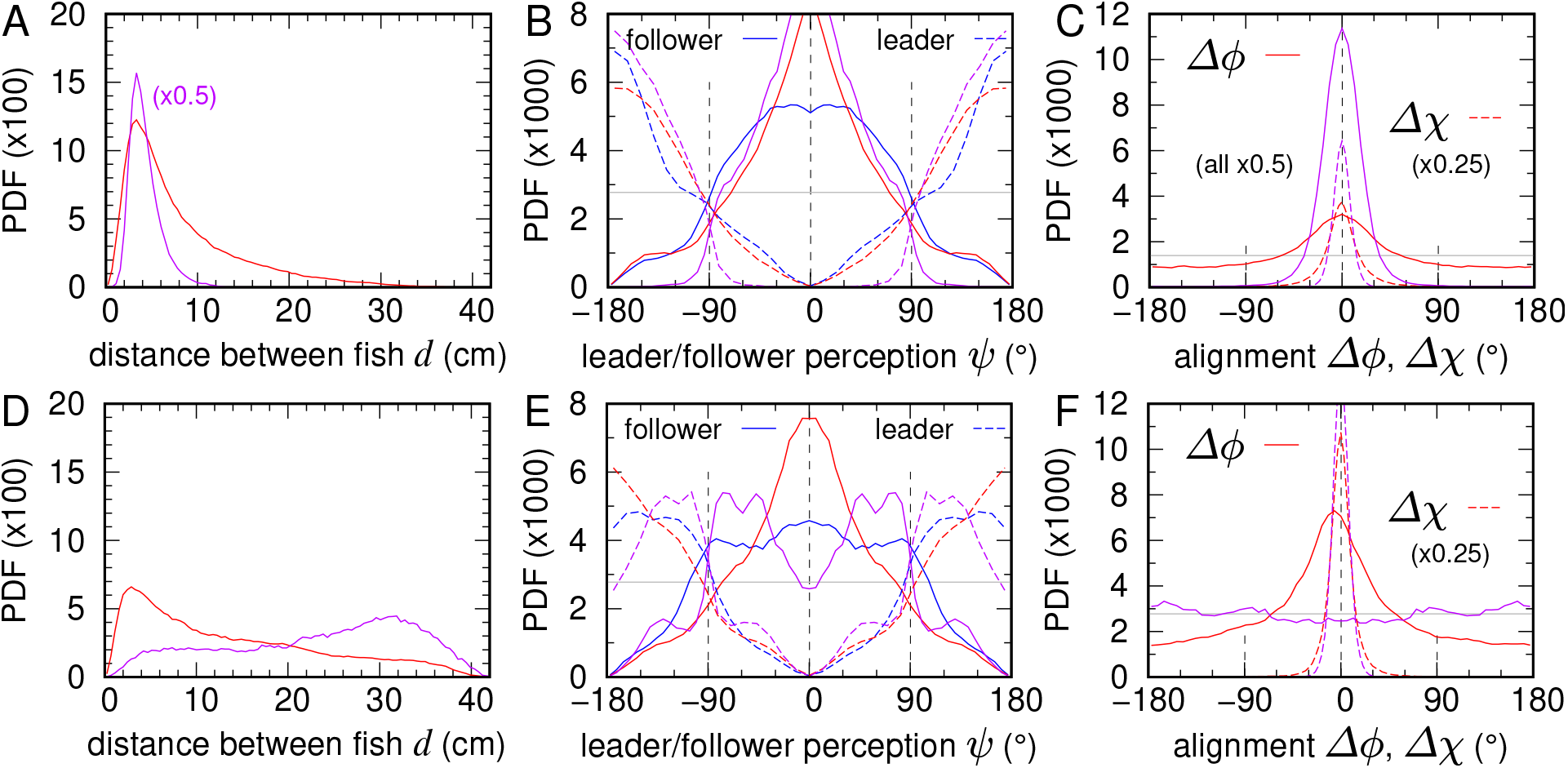
Effects of reciprocal vs. unilateral interaction with a faster virtual partner on collective coordination. Probability density functions (PDFs) of collective variables measured for pairs composed of a real fish (red) and a faster virtual fish (blue), controlled by the 2R-based model, compared with numerical simulations where both agents behave like the virtual one (violet). The mean speed of the virtual fish is increased to v 10 cm/s. (**A–C**) Condition C2: bidirectional interaction, where the virtual fish responds to the real one. (**D–F**) Condition C3: unilateral interaction, where the virtual fish behaves independently of the real one. (**A, D**) Horizontal inter-individual distance *d*. (**B, E**) Viewing angle *Ψ* for geometrical leader and follower. (**C, F**) Heading alignment between the real and virtual fish in the horizontal plane Δ*ϕ* and vertical direction Δ*χ*.

As expected, in both C2 and C3 the real fish swims faster than in the 2R baseline, but not substantially faster than in C1. The speed distributions peak at v_C2_ ≈ 7.5 cm/s (Fig. 5A) and v_C3_ ≈ 9 cm/s (Fig. 5D), close to v_C1_ ≈ 8 cm/s (Fig. 4A). However, high-speed excursions (v > 15 cm/s) are more frequent in both C2 and C3 than in C1.

In C2, the virtual fish exhibits nearly identical speed and depth PDFs as the real fish (Fig. 5A and SI Appendix Fig. S6A), showing that both fish successfully track each other. Their distance to the wall *r*_w_ and their wall-incidence angle *θ*_w_ converge towards values observed for two real fish (Figs. 5B–C). These results indicate that the higher speed of the virtual fish enhances mutual responsiveness and improves bidirectional coordination compared with C1.

From the collective perspective, the distance between fish is considerably smaller in C2 (Fig. 6A) than in C1 (Fig. 4G), and the distribution of heading differences (Δ*ϕ* and Δ*χ*) is much more peaked around zero (Fig. 6C), demonstrating higher alignment (see also SI Appendix, Movie S2 and SI Appendix, Fig. S2). This suggests that the speed of the virtual fish modulates the intensity of alignment interactions. When the virtual fish swims faster, alignment between the two fish is higher. Moreover, the mean time separating a U-turn performed by one fish and that of its companion is smaller in C2 than in C1 (2.4 ± 3.0 s) and close to the 2R condition (SI Appendix, Fig. S5).

In sharp contrast, when the virtual fish ignores the real one (Condition C3), the virtual agent behaves exactly as in the null model, but the real fish still reacts socially (see also SI Appendix, Movie S3 and SI Appendix, Fig. S1D). As a result, the distance between fish is significantly larger than in C2, yet remains much smaller than expected from the null model in the absence of social interactions (Fig. 6D). The alignment between the real and virtual fish, although reduced relative to C1, is still more pronounced than in C2 (Fig. 6F), meaning that the alignment between fish is higher when interactions are unidirectional.

Moreover, the unilateral social response in C3 causes the real fish to behave predominantly as a follower, doing so for 57.2% of the time, compared to 38.8% in C1 and 46.6% in C2. In these periods, The real fish perceives the virtual partner as swimming ahead (|*Ψ*| < 90^°^; solid red line in Fig. 6E). This persistent follower role indicates that the real fish maintains coordinated swimming even when the virtual fish provides no reciprocal social feedback.

## 3 Discussion

Understanding how real organisms coordinate their movements requires models that accurately capture the rules they follow.^2, 35^ A powerful way to validate such models is to directly insert a simulated organism into a natural interaction and observe how a real animal responds.^14, 13, 32, 16, 23, 17^ In this study, we mixed a real fish with its digital twin by using a closed-loop virtual reality system. We first measured and reconstructed the social interaction functions between two real fish swimming freely in a semi-spherical bowl, allowing precise quantification of their three-dimensional trajectories and mutual responses. Then, we developed a 3D behavioral model incorporating these functions and demonstrated that it reproduced natural coordination dynamics. Finally, we embedded this model in an immersive VR setup so that a virtual fish could interact in real time with a real fish. We used these experiments to test whether the model captures the social forces that govern pairwise swimming. This study provides a crucial step toward strong validation of models of collective behavior in living systems.

A first result is that the reconstructed interaction model quantitatively reproduces both individual and collective level dynamics in 3D simulations involving two virtual fish. When the agents are controlled by the interaction functions extracted from real pairs, they exhibit key features of natural swimming behavior: they remain close to the wall, maintain a preferred depth, and sustain an attraction that is strong enough to maintain tight spatial cohesion. Their distributions of speed, angular velocity, and inter-individual positioning closely match experimental baselines involving two real fish. These results are consistent with prior 2D reconstructions of attraction and alignment forces in burst-and-coast swimmers^5, 34^ and support the general idea that visual cues alone can sustain coordinated motion in social fish.^36, 29, 30, 31^ These results also demonstrate that the 3D self-propelled particle model captures the core geometric and sensory constraints shaping fish interactions, and that data-driven interaction rules can generate realistic collective motion in 3D environments.

When we allowed the virtual fish to interact in real time with a live fish (Condition C1), some differences emerged. The real fish often swam faster and executed sharper turns than in natural pairs, leading to greater separation and reduced alignment. Although the interaction rules were correct and allowed the real and virtual fish to coordinate their swimming when they were close to each other, the virtual partner could not always match the rapid acceleration and tight turning exhibited by a real fish. As a result, mutual responsiveness was weakened and insufficient to fully maintain cohesion. These results reveal a key requirement for model realism, showing that virtual agents must not only follow accurate perceptual and social rules but also reproduce biomechanical limits and actuation capabilities^37^ in order to sustain stable coordination.

The second important finding is that swimming speed modulates the strength of social coordination. When we increased the preferred speed of the virtual fish (Condition C2), the real fish matched that speed almost exactly. Both individuals successfully tracked each other. Their distance from the wall and their alignment angles became close to those of two real fish. Spatial cohesion improved markedly compared to Condition C1. These results demonstrate that a faster virtual partner improves joint coordination. The real fish intensified its social response when swimming with a faster conspecific. When movement becomes faster, it amplifies the salience of social cues^38^ and elicits stronger reciprocal feedback. Similar observations have been made in studies of flocking birds or schooling fish exposed to threats,^24, 25^ where speed elevation leads to stronger alignment and reduced reaction delays. Consistent with these findings, experiments using an extremely social robotic fish have shown that an individual’s mean swimming speed strongly influences group cohesion, polarization, and spatial structure.^26^ Our closed-loop experiments provide a controlled demonstration that kinematic context can push the system toward more stable collective motion.

A third key result concerns unilateral interactions (Condition C3). Even when the virtual fish no longer responded to the real one, the pair maintained a striking level of coordination. The real fish continued to match the virtual partner’s heading and orientation, far exceeding expectations from a null model lacking social feedback. The main difference was an increase in separation distance. Because the virtual agent never adjusts its trajectory to preserve proximity, unlike a responsive partner, brief episodes of larger spacing accumulate over time. This shift produces a distance distribution skewed toward higher values, not because the real fish reduces coordination, but because the interaction becomes inherently non-reciprocal. The strong social persistence maintained by the real fish highlights that the behavioral strategy is sufficiently robust even when reciprocal information is degraded. This finding sheds light on interaction asymmetries and leadership dynamics in collective systems.^27, 24, 28, 39^ Recent volumetric VR experiments with freely swimming zebrafish further show that coordination involves not only spatial responses but also a robust, out-of-phase temporal alternation of burst events that emerges exclusively when feedback is reciprocal.^17^ This temporal coupling enhances an individual’s ability to track sudden directional changes in a partner, revealing a complementary mechanism of coordination that sits alongside the spatial and kinematic factors explored in our closed-loop experiments. The comparison highlights that reciprocity governs not only the geometry of coordination but also its fine-scale temporal structure, pointing to a richer interplay between timing, perception, and social responsiveness in fish.

Every experimental system has limitations. The virtual fish cannot match the full biomechanical capabilities of a real one. In particular, the model does not account for hydrodynamic effects that real fish can perceive at close range and use to coordinate their motion through neighbor-induced flows, such as alternating vortex patterns that can be exploited to reduce energetic cost or enhance propulsion.^40, 41^ The degree to which real fish perceive the projected stimulus as a mate may also vary with context.^23^ Moreover, pair interactions represent only the simplest unit of schooling dynamics, while real groups involve multiple neighbors, distributed sensing, and rapid switching of social roles. These constraints likely contributed to reduced alignment in VR conditions. Despite these constraints, our results demonstrate that closed-loop interactions in virtual reality can produce highly realistic social responses. This approach offers new opportunities to test mechanisms that are difficult to isolate in group experiments.

Overall, our findings demonstrate that real fish can form coordinated pairs with model-driven virtual conspecifics, confirming that the social interaction functions reconstructed from real pairs reliably reproduce natural collective behavior. Coordination increases with swimming speed and remains robust even when the model is challenged by unilateral feedback. These findings establish that mixing real and simulated agents is not only feasible but yields biologically meaningful interactions that directly validate the mechanisms encoded in behavioral models.

By placing a digital twin into a natural feedback loop, we overcome a common limitation of collective behavior models and move beyond validation based solely on statistical agreement to directly testing behavioral responses.^42, 43, 44, 6^ Our approach tests whether a model elicits the correct actions in the real organism itself, a far stronger and more mechanistic benchmark.

Several future improvements are already tractable. Enhancing the biomechanical realism of the virtual agent and scaling to multi-agent interactions will allow us to test complex hypotheses about perception, the combination of interactions between neighbors, and decision-making in collective behavior. Similar closed-loop VR methods have begun to reveal prediction error signals in zebrafish^15^ and speed-dependent modulation of coordination in fish groups,^36, 26^ highlighting the relevance of our approach to sensory and cognitive dynamics.

More broadly, this hybrid architecture offers a versatile platform for hypothesis testing in living systems. By merging data-driven models with interactive VR, we create adaptive, testable digital twins for collective behavior. This development paves the way toward mixed swarms of biological and digital agents, enabling experiments that would be impossible using only real or simulated organisms. Similar hybrid approaches have successfully merged robotic and animal societies,^45, 46, 47^ while interactive VR platforms now allow real-time bidirectional feedback between real and virtual agents.^16, 23^ Our work therefore represents a critical step toward fully integrative, closed-loop studies of perception, decision-making, and collective intelligence in animal groups.^48, 49^

## 4 Materials and methods

### Ethics statement

Experiments were approved by the Animal Experimentation Ethics Committee C2EA-01 of the Toulouse Biology Research Federation and were performed in an approved fish facility (A3155501) under permit APAFIS#27303-2020090219529069 v8 in agreement with the French legislation. All procedures were designed to minimize stress and handling. Fish were transferred from rearing tanks to the experimental setup with minimal manipulation. Each individual was used in only one one-hour experimental session per week. Swimming ability was monitored throughout; fish exhibiting impaired or absent swimming activity were excluded and replaced. No animals were sacrificed during this study.

### Study species

Rummy-nose tetras (*Hemigrammus rhodostomus*) were purchased from Amazonie Labège in Toulouse, France. Fish were kept in 16 L aquariums on a 12:12 hour, dark:light photoperiod, at 24.9^°^ C (±0.8^°^ C) and were fed *ad libitum* with fish flakes. The average body length of the fish used in these experiments is 3.1 cm.

### Experimental setup

We used a closed-loop virtual reality system specifically developed to study real-time interactions between freely swimming fish and computer-generated virtual conspecifics.^23^ The setup consisted of a hemispherical acrylic bowl (diameter 52.2 cm, depth 14.6 cm, filled with 15 L of water) placed above a DLP LED projector that displayed anamorphically rendered virtual fish onto the bowl wall. Fish movements were tracked from above using an Intel RealSense D435 depth camera positioned 47.9 cm above the water surface. Infrared illumination (8 IR lamps, 850 nm) enabled robust tracking under controlled light conditions. A dedicated workstation (Dell Precision 3640 with NVIDIA RTX 3070) processed all tasks in real time, including 3D tracking, trajectory simulation of virtual fish, and rendering.

The system included three software modules: a Python-based 3D tracking pipeline, a C++ trajectory simulator, and a Unity rendering engine. In this study, the system was used in closed-loop mode: the position and heading of the virtual fish were continuously controlled by the behavioral model parameterized for rummy-nose tetras, exploiting the current position and speed of the real fish. Real and virtual fish positions were exchanged via UDP at 30–90 Hz, ensuring precise, biologically relevant interactions. The same setup was also employed to study the behavior and interactions of pairs of real fish, using the same structured background and identical lighting conditions as in experiments involving one real and one virtual fish.

### Experimental procedure

We conducted four series of experiments combining behavioral recordings of real fish pairs and closed-loop VR assays. In the first series, we tracked pairs of *H. rhodostomus* swimming freely in a confined 3D space to characterize their spontaneous motion and social interactions. Sixteen one-hour replicates were performed. The collected trajectories provided reference values for swimming speed, spatial positioning, and alignment between individuals, and were used to extract analytical functions describing self-propulsion, wall avoidance, and social interaction terms. These functions formed the basis of the behavioral model controlling the virtual fish. In subsequent experiments, a single real fish was introduced into the hemispherical VR tank and exposed to a virtual conspecific whose motion was governed in real time by this model. The position and heading of the virtual fish were continuously updated in closed loop based on the current position and velocity of the real fish. Three experimental conditions were tested, each defined by a distinct set of model parameters (SI Appendix, Table S1). Condition C1 corresponded to the reference parameter set reproducing the dynamics of two real fish swimming together (see SI Appendix, Movie S1). Condition C2 used higher self-propulsion parameters to match the increased swimming speed observed in C1 (see SI Appendix, Movie S2). Condition C3 used the same parameters as C2 but with all social interaction terms disabled, such that the virtual fish moved independently of the real one (see SI Appendix, Movie S1). Eight one-hour replicates were performed under C1 and C3, and twelve under C2. Each trial began after a 10 min acclimation period.

### Preprocessing of fish trajectories

Before analyzing the trajectories extracted from the experiments, the data must be preprocessed to ensure accuracy and to retain only periods in which fish are actively swimming. Time intervals during which a fish remains immobile for at least 4 seconds are discarded, as well as segments in which its instantaneous speed exceeds 25 cm/s or when it swims in a straight line at approximately constant speed for more than one second. These empirical thresholds were determined from the statistical properties of spontaneous swimming behavior and correspond, respectively, to periods of inactivity and to tracking artifacts such as sudden position jumps.

The dataset is then filtered to remove any frames in which the reconstructed position lies outside the bowl. To recover continuous temporal trajectories, gaps created during this filtering procedure are linearly interpolated whenever the duration of missing data is shorter than one second. For longer gaps, reliable reconstruction is not possible, and the trajectory is left discontinuous at that point.

The resulting trajectories therefore consist of multiple consecutive temporal segments. Each segment is subsequently smoothed by a Gaussian kernel convolution in order to reduce tracking noise while preserving the essential kinematic features of the motion,

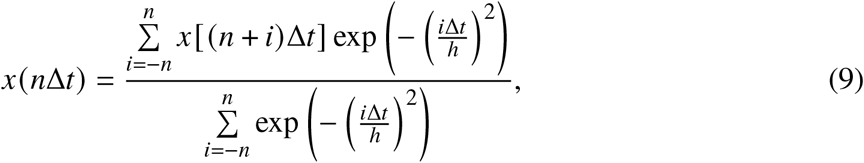

where *n* = 4*h* / Δ*t*, such that the exponential kernel weight is negligible at *i* = *n*. The kernel width is set to *h* = 0.5 s (on par with the duration of a kick arising from the burst-and-coast swimming mode of *H. rhodostomus*), which provides minimal smoothing while reducing high-frequency noise.

### Quantification of individual and collective behavior in pair of fish

Fish predominantly swim in the horizontal plane at a preferred comfort depth. Throughout the manuscript, three-dimensional quantities are denoted with a superscript 3D, horizontal (2D) quantities without a superscript, and vertical components with a subscript *z*.

The experimental bowl is modeled as the lower cap of a sphere of radius *R*_0_ = 26.19 cm, filled with 15 liters of water, resulting in a central depth of *h*_water_ = 14.73 cm. The coordinate origin is placed at the water surface above the center of the bowl, with the *z*-axis oriented downward so that depth values are positive and increase with distance from the surface. At depth *z*, the radius of the corresponding horizontal cross-section is

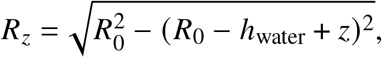

which vanishes at *z* = *h*_water_, as expected.

The fish position and velocity are given by the vectors 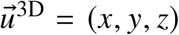 and 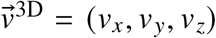. We define the distance to the bowl’s vertical axis as 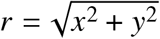 and the distance to the horizontal boundary at depth *z* as *r*_w_ = *R*_*z*_ − *r*. For consistency with previous work, we refer to this horizontal boundary as the *wall*.

The horizontal velocity is 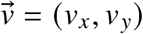, and the full 3D speed and its horizontal component are 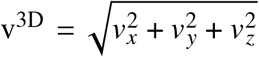 and 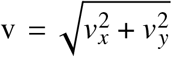, respectively. The fish heading in the horizontal plane is defined as *ϕ* = atan2 (*v*_*y*_, *v*_*x*_), ranging from −*π* to *π*, while the elevation angle is *χ* = atan (*v*_*z*_ / v), ranging in (−*π* / 2, *π* / 2) when v > 0. Heading is used to compute the angle of incidence to the wall, *θ*_w_ = *ϕ* − *θ*, where *θ* = atan2 (*y, x*) is the positional azimuth. By convention, positive headings are counterclockwise, and positive (negative) elevation indicates upward (downward) motion.

When swimming with a conspecific, the relative configuration is defined by their spatial separation and angular alignment. For a pair of fish *i* and *j*, the 3D distance is 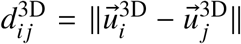 the horizontal distance 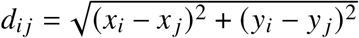, and the vertical distance 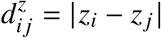. The *viewing angle Ψ*_*i j*_, defined as the angular correction required for fish *i* to orient toward fish *j*, is computed as *Ψ*_*i j*_ = atan2 (*y* _*j*_ − *y*_*i*_, *x* _*j*_ − *x*_*i*_) −*ϕ*_*i*_. Note that, in general, *Ψ*_*i j*_ *≠* − *Ψ* _*ji*_, meaning interaction geometry is not necessarily reciprocal.

We define the *geometrical leader* as the individual with the larger perception angle magnitude |*Ψ*_*i j*_|, while the other fish is considered the *geometrical follower*. Alignment differences in the horizontal and vertical planes are given by *ϕ*_*i j*_ = *ϕ* _*j*_ −*ϕ*_*i*_ = −*ϕ* _*ji*_ and *χ*_*i j*_ = *χ*_*j*_ − *χ*_*i*_ = *χ*_*ji*_.

Since only dyads are studied here, notation is simplified by omitting indices when no ambiguity arises, writing *d, Ψ*, Δ*ϕ*, Δ*χ* instead of *d*_*i j*_, *Ψ*_*i j*_, *ϕ*_*i j*_, *χ*_*i j*_.

### 3D self-propelled particle model for fish swimming in a bowl

The horizontal velocity of a fish is decomposed into components parallel and perpendicular to its instantaneous heading. Defining the unit vectors 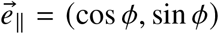 and 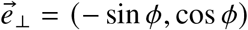, the velocity in the horizontal plane is expressed as 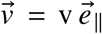. The horizontal acceleration is written as 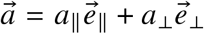, where *a*_∥_ and *a*_⊥_ are the longitudinal (speed-changing) and lateral (turning) acceleration components, respectively. Together with the vertical acceleration *a*_*z*_, these variables fully determine the temporal evolution of fish position.

Expressing each force as the product of a scalar with the unit vector of its direction, the components of fish’s acceleration in the horizontal plane are as follows:

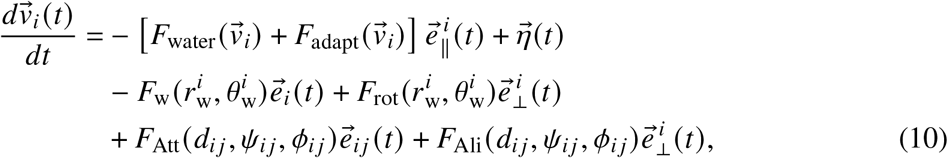

where 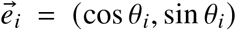, with *θ*_*i*_ = atan2(*y*_*i*_, *x*_*i*_), is the position angle of fish *i* in the horizontal plane and is perpendicular to the wall closest to the fish in the horizontal plane. 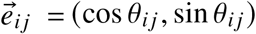, with *θ*_*i j*_ = atan2(*y* _*j*_ − *y*_*i*_, *x* _*j*_ − *x*_*i*_), is oriented along the direction from *i* to *j* in the horizontal plane. Using *Ψ*_*i j*_ = *θ*_*i j*_ − *ϕ*_*i*_ and 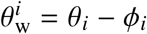, the vectors 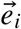 and 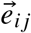 can be expressed in the fish frame as 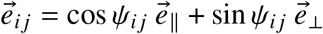 and 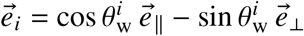.

In the intrinsic frame of fish *i*, this gives the scalar components of the acceleration:

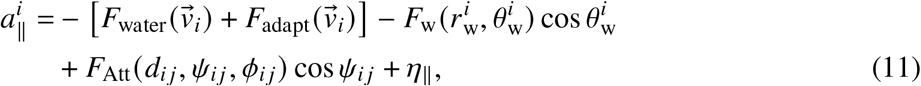

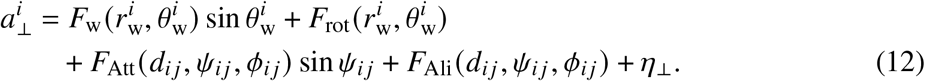

Using explicitly the decomposition of the interactions into products of single-variable functions, the final system reads

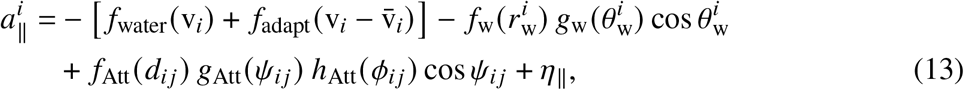

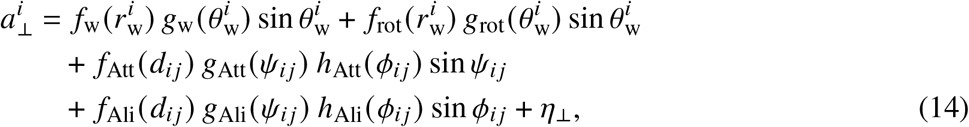

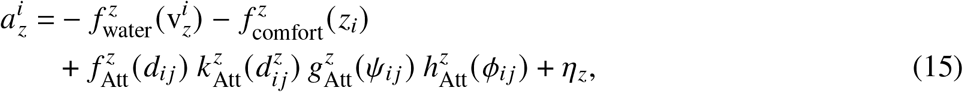

where 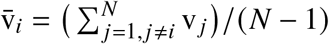 is the average speed of the neighbor(s) of *i*, which for *N* = 2 fish becomes 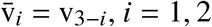.

The analytical expressions of the different functions arising in these equations of motion (5 polynomial expansions, 8 Fourier series, and 5 simple fits of tabulated functions), used in the reconstruction procedure and implemented in the model, are given in Supplementary Materials.

### Stochasticity

Behavioral fluctuations are modeled as autocorrelated noise processes in the parallel, perpendicular, and vertical directions. Noise components are defined in the fish-centered reference frame and converted into Cartesian coordinates via

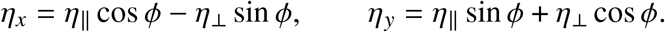

They evolve according to a discrete Ornstein-Uhlenbeck process 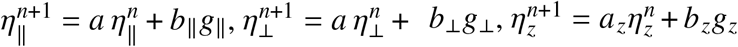 with time-step index *n*. The random variables *g*_∥_,*g*_⊥_,*g*_*z*_ are independent standard normal deviates generated using 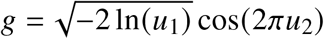, with *u*_1_, *u*_2_ ∼ 𝒰 (0, 1) (Box-Muller transform). The parameters corresponding to the Ornstein-Uhlenbeck dynamics are given by

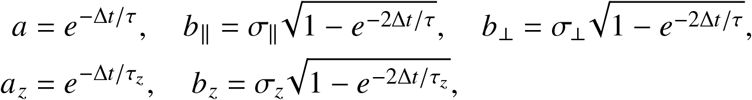

where *σ*_∥_, *σ*_⊥_, and *σ*_*z*_ set noise amplitudes, and *τ, τ*_*z*_ are correlation times in the horizontal and vertical directions, respectively, that we took equal in practice. The occurrence of finite correlation times for the noises is consistent with the true burst-and-coast nature of the fish swimming mode, where a fish alternates a very brief acceleration period (of typical duration 0.1 s) with a passive gliding period lasting typically 0.5 s, where the fish barely changes its heading.

### Boundary rejection rules

In this agent-based model, (*x, y, z*) denotes the center of mass of the fish. Since fish have a finite body length, the predicted position at step *n* + 1 may place the head outside the water volume. Such moves are rejected.

The position of the head is obtained from

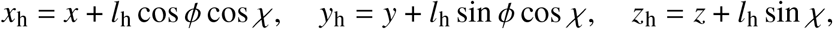

where *l*_h_ is the head-center distance.

Two rejection cases are implemented:

- **Collision with the wall:** If the head lies beyond the wall surface, 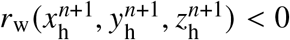, the previous position (*x*^*n*^, *y*^*n*^, *z*^*n*^) is restored, and the fish is reoriented to swim nearly tangentially to the wall (adding a small random angle of order *B* ≈ 0.15 radian):

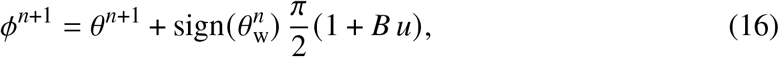

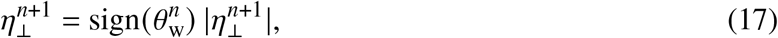

where *u* ∼ 𝒰 (0, 1). This ensures 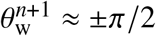, i.e., near alignment with the wall.
- **Exiting the water surface:** In the very rarely observed cases where the head is predicted to lie above the surface or below the bowl bottom, the depth is restored and the vertical speed is reversed: *z*^*n*+1^ = *z*^*n*^ and 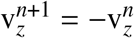.

## Acknowledgments

None.

## Supplementary Materials

## Supplementary Text

### Analytical expressions of the extracted social interaction functions

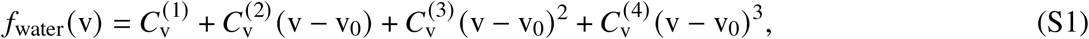

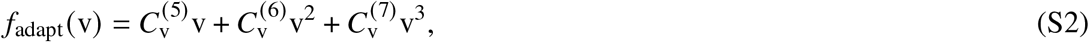

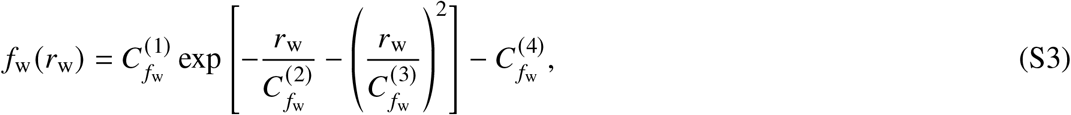

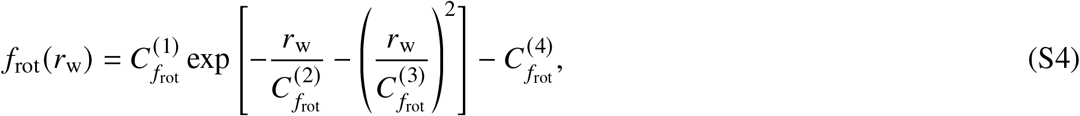

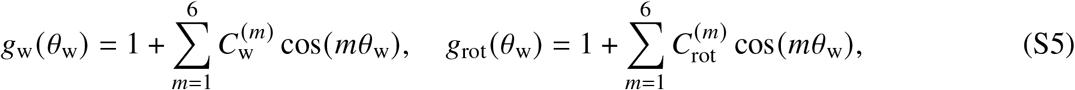

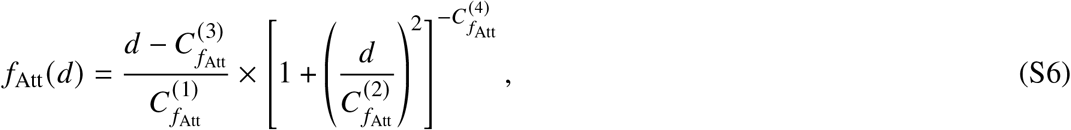

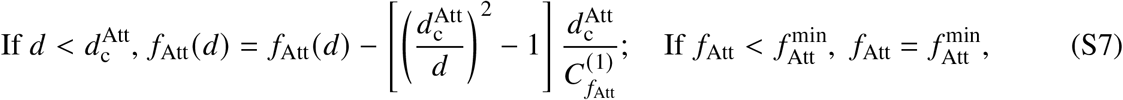

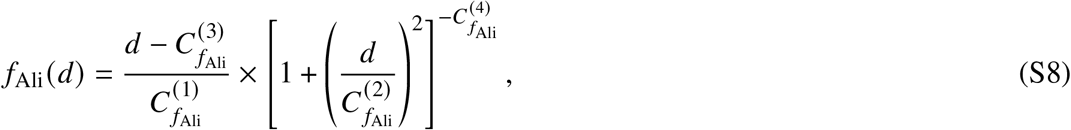

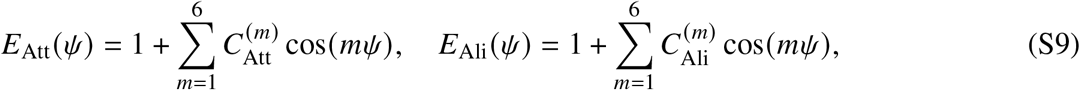

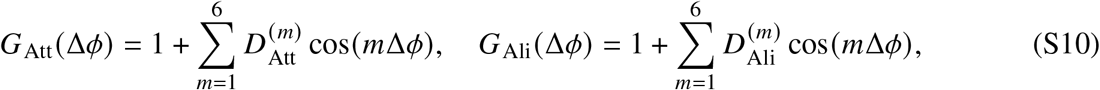

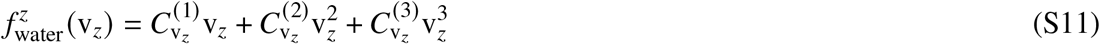

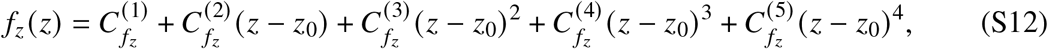

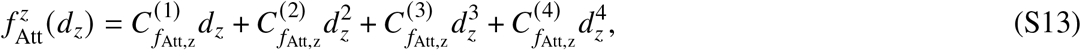

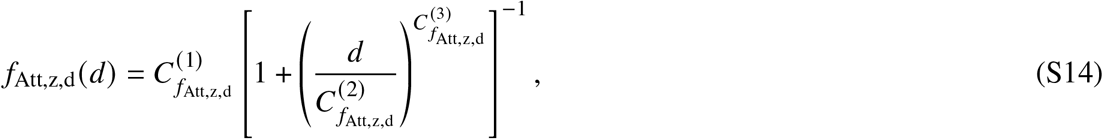

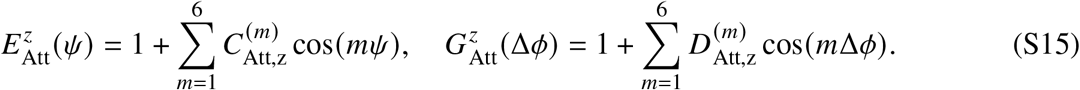

Values of parameters and coefficients are given in Table S1.

## Supplementary Tables

**Table S1:** Values of model parameters used in conditions C1, C2 and C3. The values of simulation parameters used in Condition C1 are those that provide the best match to experiments with two real fish.

| Symbol | Parameter | Condition C1 (2 real fish) | C2 and C3 |
| --- | --- | --- | --- |
| $\sigma_{\parallel}$ | Noise parallel component | 3.4 | 3.3 |
| $\sigma_{\perp}$ | Noise perpendicular component | 2.9 | 2.8 |
| $\sigma_z$ | Noise vertical component | 2.9 | 2.35 |
| $\tau_{XY}, \tau_Z$ | Correlation times (s) | 0.25 | 0.25 |
| $C_{v_0}$ | Comfort speed (cm/s) | 7.2 | 12 |
| $D_{f_{Att}}$ | Attraction equilibrium distance | 30 | 40 |
| $C_{water}$ | Friction modulation factor | 1 | 0.5 |
| $C_{water}^z$ | Vertical friction factor | 1 | 0.75 |
| $C_{f_w}$ | Wall repulsion modulation factor | 0.6 | 3.9 |
| $C_{f_{Att}}$ | Attraction force factor | 2 | 3.5 |
| $C_{f_{Ali}}$ | Alignment force factor | 1.4 | 2 |
| $C_{f_{Att}}^z$ | Vertical attraction factor | 1 | 2 |

**Figure S1:**
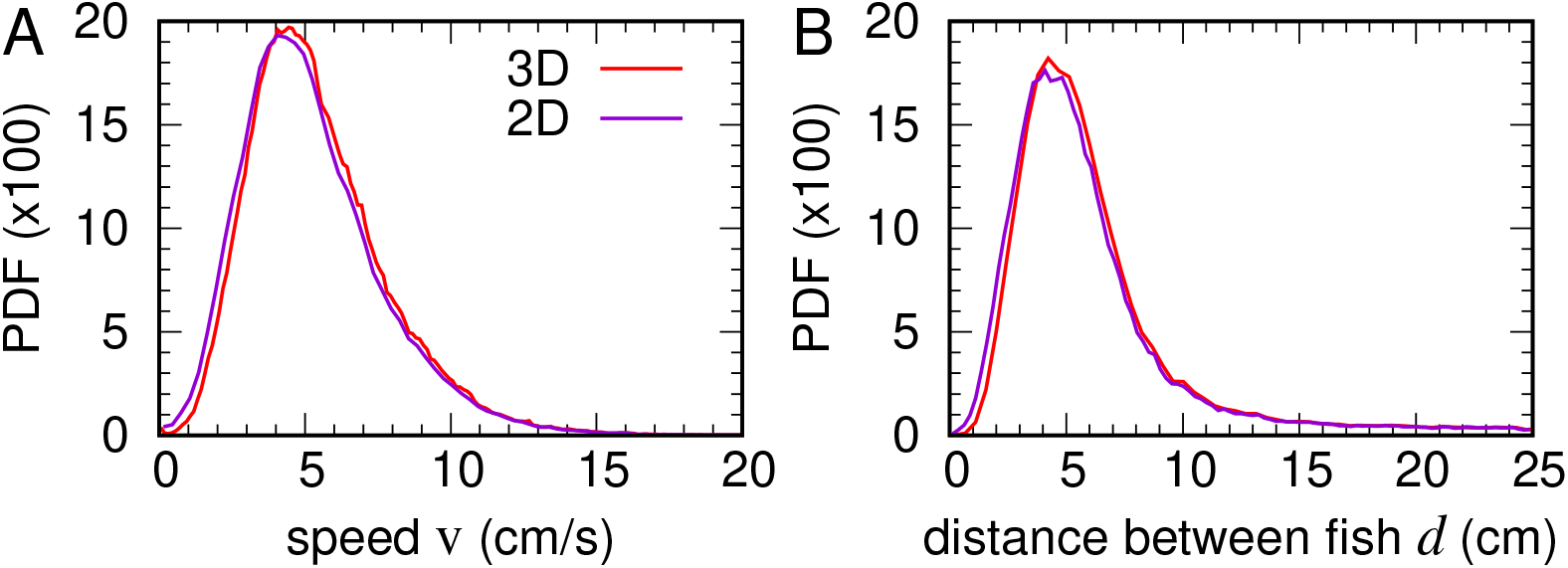
Comparison of three-dimensional speed and distance between fish with their projections on the horizontal plane. (**A**) Probability density functions (PDF) of the 3D-speed (red) and of its projection on the horizontal plane (violet). (**B**) PDFs of the 3D-distance between fish (red) and of its projection on the horizontal plane (violet).

**Figure S2:**
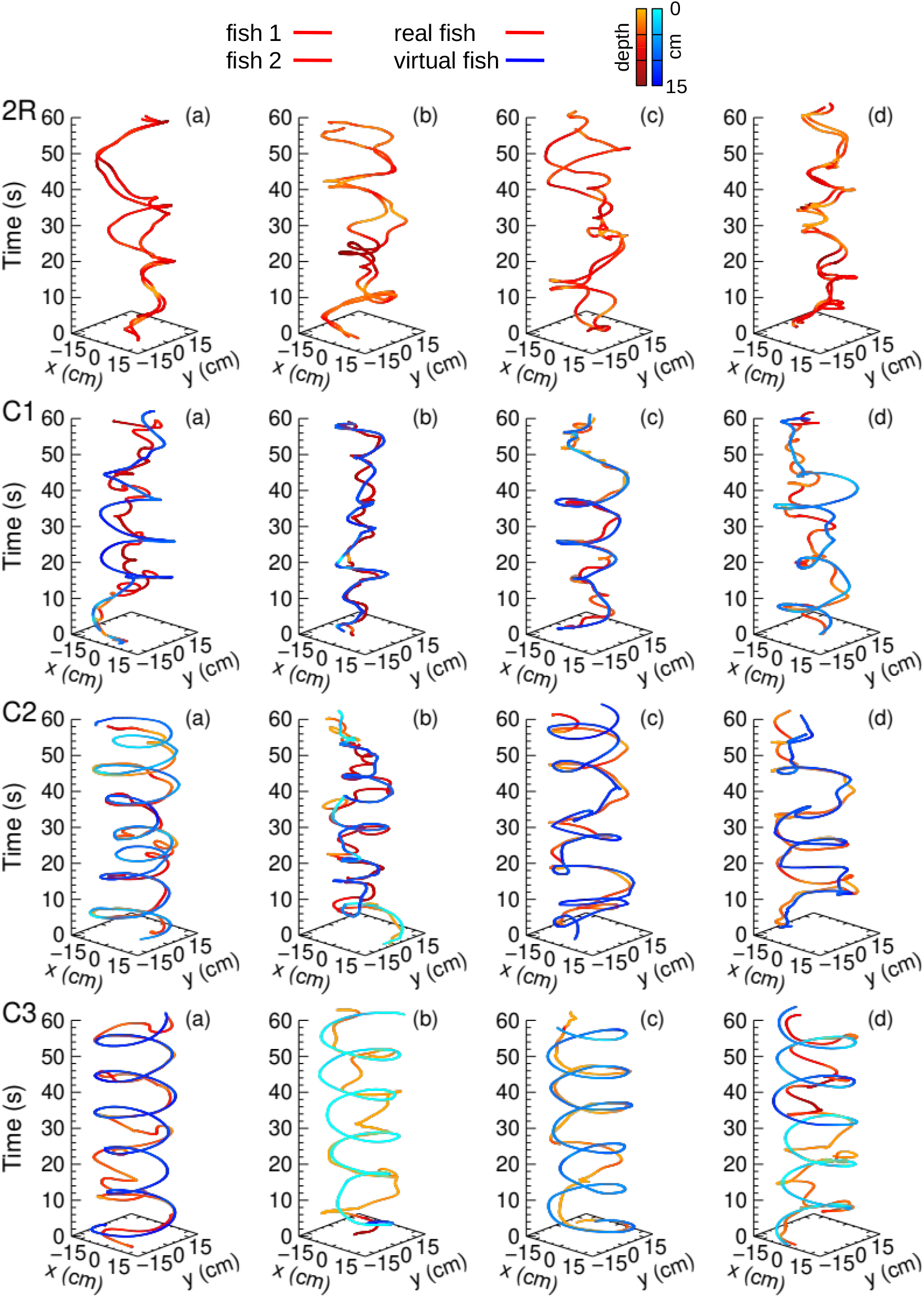
Real and virtual fish trajectories. (**2R**) Four representative trajectories of a pair of *Hemigrammus rhodostomus* swimming freely in a hemispherical bowl for one minute. (**C1**) Four representative trajectories of a real fish (red) and a virtual fish (blue) that reproduces the behavior of a real fish swimming in a pair for one minute in Condition C1 (v ≈ 5 cm s^−1^). (**C2**) Same as C1 but with the virtual fish moving at twice the natural speed (v ≈ 10 cm s^−1^) and with reciprocal interactions. (**C3**) Same as C2 but without reciprocal interactions.

**Figure S3:**
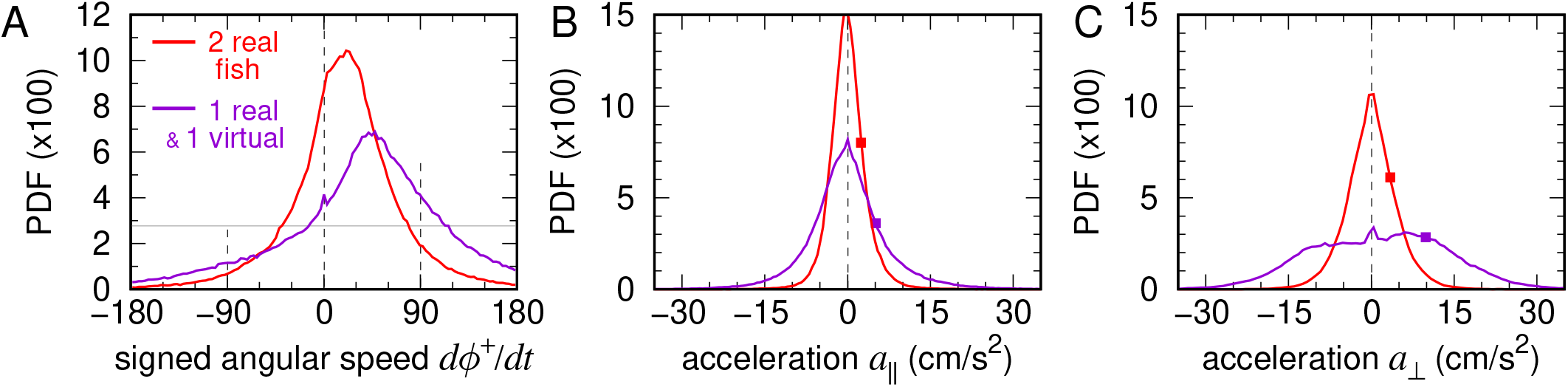
Real–virtual interactions induce stronger turning accelerations. Probability density functions (PDFs) of kinematic variables describing turning maneuvers when the focal fish interacts with a real (red) or a virtual (violet) partner: (**A**) Signed angular velocity *dϕ*^+^/*dt* (sign defined by the incidence angle *θ*_w_), (**B**) Parallel acceleration *a*_∥_, (**C**) Perpendicular acceleration *a*_⊥_. Squares in (**B**,**C**) indicate the absolute mean accelerations ⟨|*a*_∥_|⟩ and ⟨|*a*_⊥_|⟩, showing that perpendicular accelerations are more than twice as large as parallel ones when swimming with a virtual partner.

**Figure S4:**
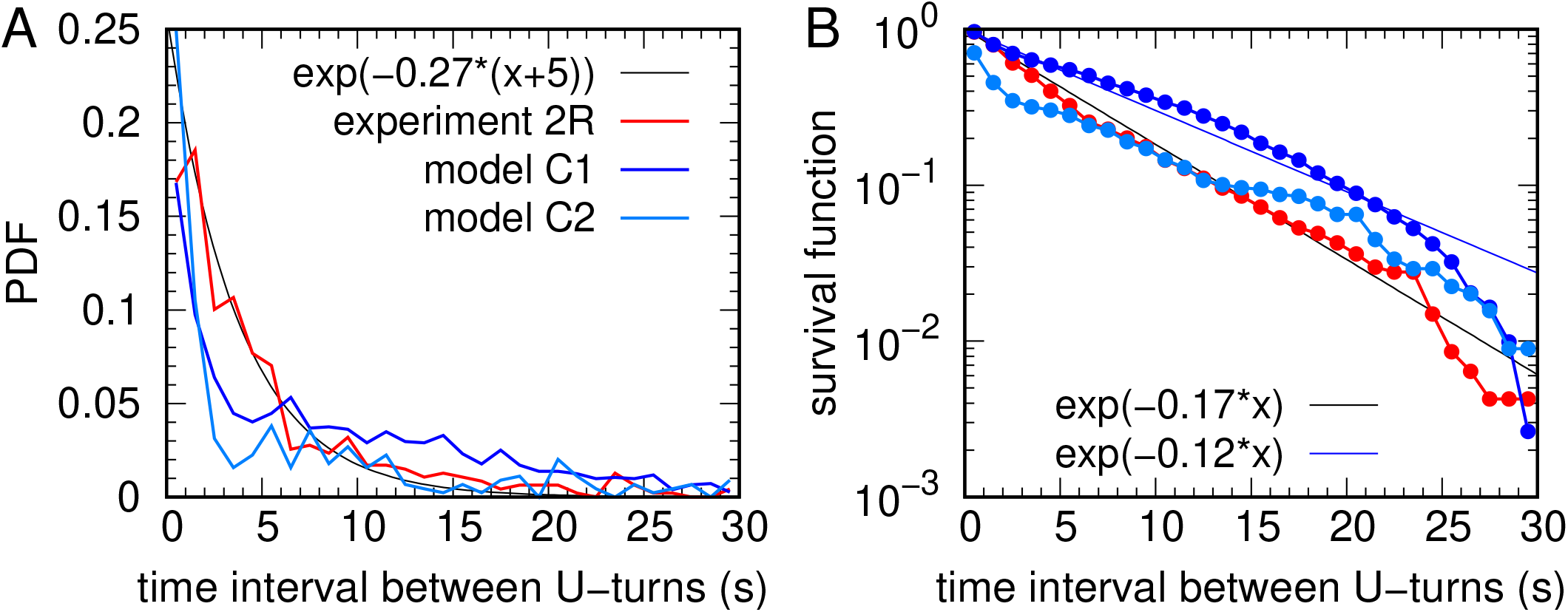
Time interval between U-turns. (**A**) Probability density function (PDF) of the time interval between two consecutive U-turns (angle change >100^◦^ in 0.5 s) performed by the same fish, measured in experiments with two real fish (red), and in numerical simulations of the model with parameters corresponding to Conditions C1 (dark blue) and C2 (light blue). (**B**) Corresponding survival curves. Straight lines indicate exponential fits.

**Figure S5:**
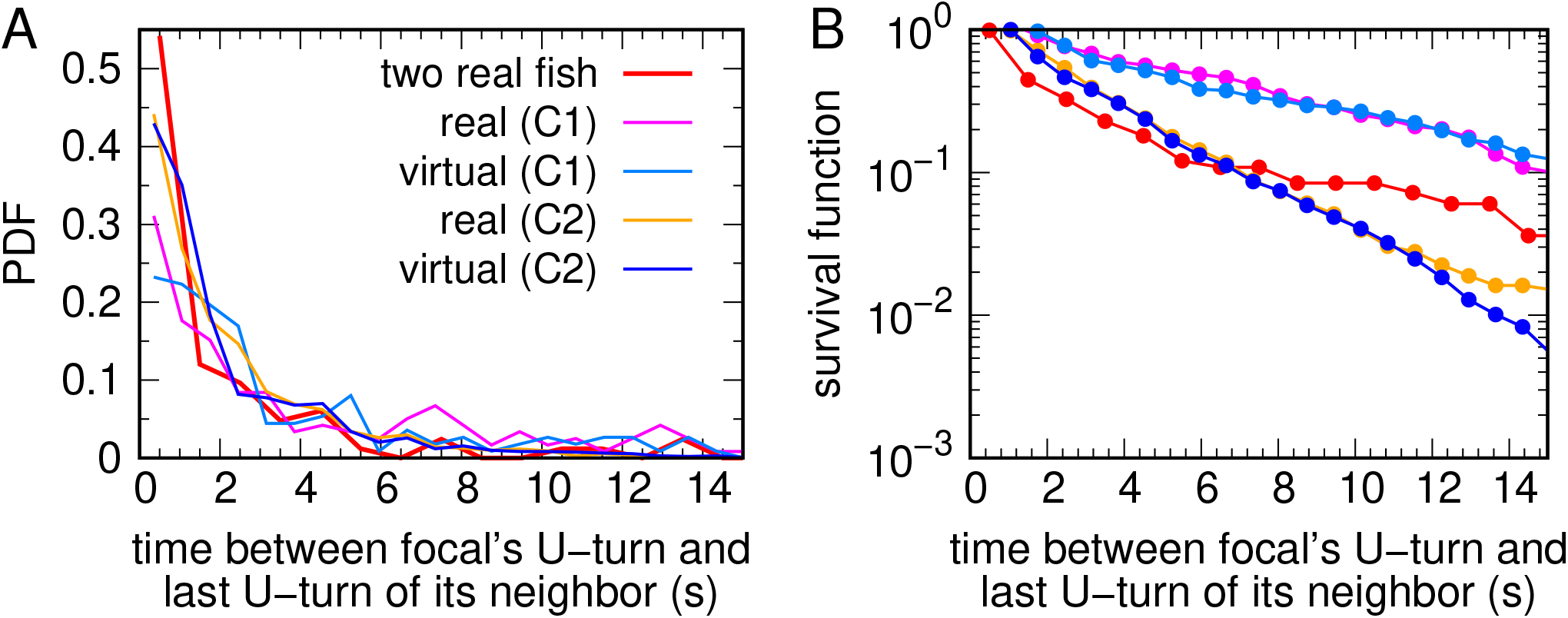
Time interval between a U-turn of the focal fish and the U-turn of its neighbor. (A) Probability density function (PDF) of the time interval between between a U-turn of the focal fish and the U-turn of its neighbor (angle change >100^◦^ in 0.5 s), measured in experiments with two real fish (red), and in Conditions C1 and C2. (B) Corresponding survival curves.

**Figure S6:**
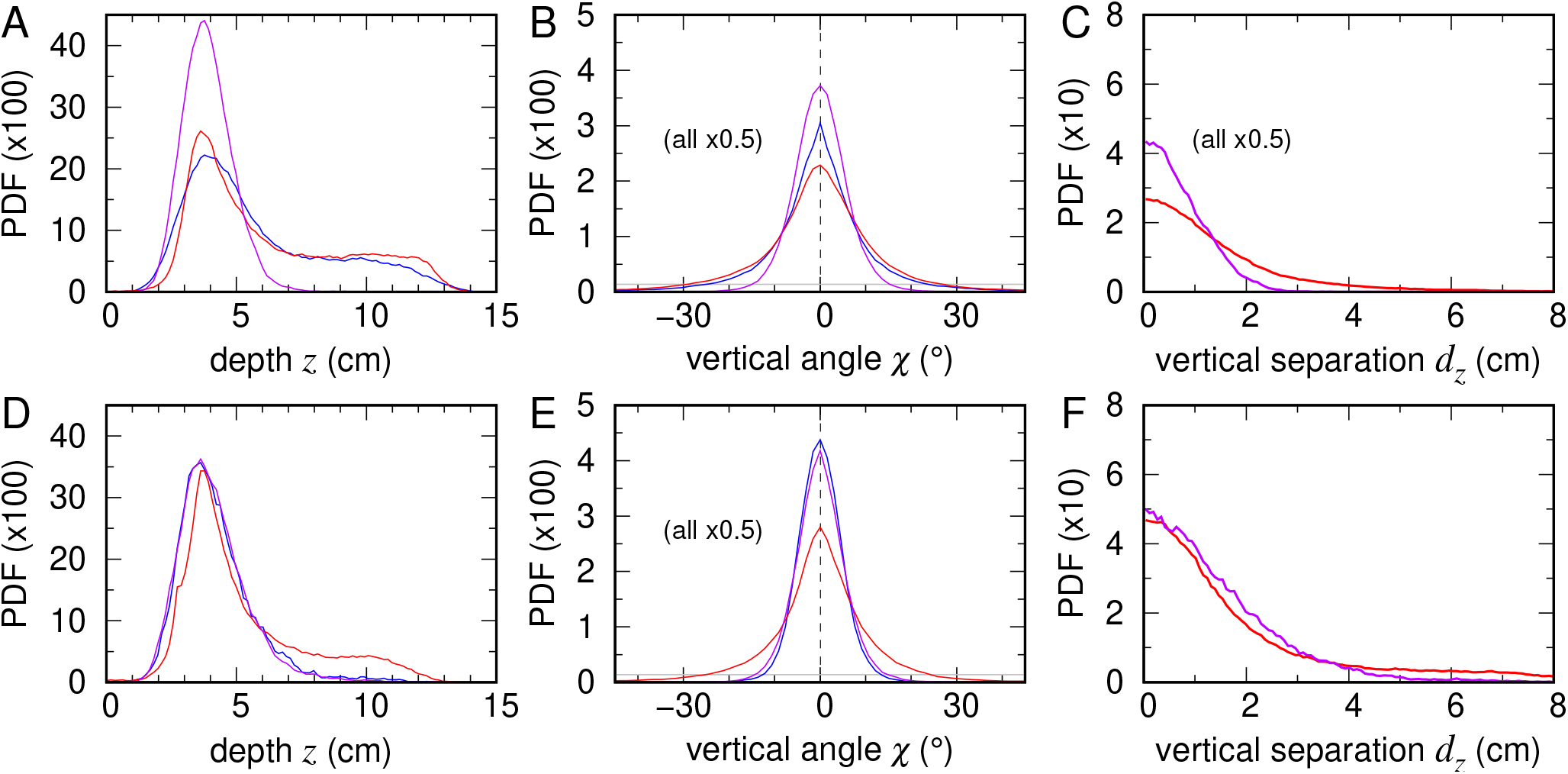
Effects of reciprocal vs. unilateral interaction with a faster virtual partner on vertical motion. Probability density functions (PDFs) of vertical behavioral variables for a real fish (red), a faster virtual fish (blue), and simulations of the 2R-based model (violet), using a higher mean virtual speed v ≈ 10 cm/s. (**A–C**) Condition C2: social interactions enabled (bidirectional coordination). (**D–F**) Condition C3: social interactions disabled (unilateral interaction). (**A, D**) Swimming depth *z*. (**B, E**) Elevation angle *χ*. (**C, F**) Vertical inter-individual distance *d*_*z*_.

## Supplementary Movies

**Movie S1. Behavioral responses of a real fish interacting with a realistic virtual partner**. Video excerpt of an experiment in Condition C1 with a real fish interacting with the anamorphic projection of a virtual conspecific whose motion and interactions with the real fish are controlled in real time by the model. In this condition, the virtual fish is a perfect clone, reproducing the behavior of a real fish in a pair with a mean speed v ≈ 5 cm/s. Top-Left: user interface allowing the real-time visualization of the trajectories of the real and virtual fish (in the xy- and xz-planes) and the instantaneous modification of the parameters of the model driving the virtual fish. Bottom-Left: real-time 3D tracking of the real fish. Right panel: anamorphic rendering of the virtual fish projected onto the bowl by the rendering application according to the 3D position of the real fish.

**Movie S2. Behavioral response of a real fish to a faster virtual conspecific with reciprocal interactions**. Video excerpt of an experiment in Condition C2 with a real fish interacting with the anamorphic projection of a virtual conspecific whose motion and interactions with the real fish are controlled in real time by the model. In this condition, the virtual fish is moving at twice the natural speed observed for real fish in condition C1 (v ≈ 10 cm/s). Top-Left: user interface allowing the real-time visualization of the trajectories of the real and virtual fish (in the xy- and xz-planes) and the instantaneous modification of the parameters of the model driving the virtual fish. Bottom-Left: real-time 3D tracking of the real fish. Right panel: anamorphic rendering of the virtual fish projected onto the bowl by the rendering application according to the 3D position of the real fish.

**Movie S3. Behavioral response of a real fish to a faster virtual conspecific with reciprocal interactions**. Video excerpt of an experiment in Condition C3 with a real fish interacting with the anamorphic projection of a virtual conspecific. In this condition, the virtual fish behaves as an isolated agent, moving at a speed of approximately v ≈ 10 cm s^−1^ according to the model’s dynamics, and does not respond to the real fish. Top-Left: user interface allowing the real-time visualization of the trajectories of the real and virtual fish (in the xy- and xz-planes) and the instantaneous modification of the parameters of the model driving the virtual fish. Bottom-Left: real-time 3D tracking of the real fish. Right panel: anamorphic rendering of the virtual fish projected onto the bowl by the rendering application according to the 3D position of the real fish.

